# The Icelandic mutation APP^A673T^ on amyloid-β plaque burden in the 5xFAD Alzheimer model

**DOI:** 10.1101/2025.11.05.686739

**Authors:** Anne Anschuetz, Renny Listyono, Thomas Vorley, Bettina Platt, Charles R. Harrington, Gernot Riedel, Karima Schwab

## Abstract

The protective Icelandic mutation in the amyloid precursor protein (APP) gene, APP^A673T^, identified in Icelandic and other Nordic populations is associated with a significantly lower risk of developing Alzheimer’s disease (AD). Conflicting results have been reported for the APP^A673T^ mutation in various knock-in models of AD, but its effect in 5x familial AD (5xFAD) mice has never been investigated. We have crossed C57Bl6/J mice expressing a single point mutation edited into the murine APP gene via CRISPR-Cas gene editing, termed APP^A673T^, with 5xFAD mice that overexpress human APP carrying the Swedish (K670N/M671L), Florida (I716V), and London (V717I) mutations as well as human presenilin-1 (PS1) with two mutations (M146L and L286V); the resulting mice were termed 5xFADxAPP^A673T^. We have investigated amyloid beta (Aβ) pathology in 5xFADxAPP^A673T^, 5xFAD and their respective controls, APP^A673T^ and C57Bl6/J wild types, at 6-months of age using immunohistochemistry, immunoblotting, and ELISAs. We found a moderate yet significant reduction for Aβ plaque size in male 5xFADxAPP^A673T^ compared to 5xFAD. No differences were observed for soluble/insoluble Aβ40 and Aβ42 levels per se, but lower plaque count/area was found in 5xFADxAPP^A673T^ when Aβ42/Aβ40 ratios were low, suggesting a genotype-dependent sensitivity to Aβ aggregation and accumulation. Therefore, the APP^A673T^ mutation has the potential to modify Aβ pathology in 5xFAD mice at the age of 6 months.

## Introduction

Alzheimer’s disease (AD) is a progressive degenerative brain disorder affecting memory, cognition and behaviour (Scott and Barrett, 2007). It is the most common cause of dementia accounting for 60-70% of cases, and numbers are expected to exceed 150 million cases worldwide by 2050 (GBD 2019 Dementia Forecasting Collaborators, 2022). Pathologically, end-stage AD is characterised by the formation of neurofibrillary tau tangles and extracellular amyloid beta-protein (Aβ) plaques (Alzheimer, 1907), and both pathologies may contribute to the neuronal dysfunction and cognitive decline observed in AD (Zhang et al., 2021). Amyloid-β is generated from the amyloid precursor protein (APP) through a series of enzymatic cleavages (Cho et al. 2022, Gabriele et al. 2022. In the amyloidogenic pathway, APP is first cleaved by β-secretase to produce a secreted form of APP (sAPPβ) and a membrane-bound carboxyl terminal fragment (βCTF or C99) - the latter is further cleaved by the γ-secretase complex (a four-unit protease complex with presenilins as the catalytic subunits) to release Aβ peptides including Aβ40 and Aβ42. Both, Aβ40 and Aβ42, are neurotoxic and an increase in the Aβ42/Aβ40 ratio has been associated with a more pronounced plaque pathology due to higher oligomerization of Aβ42 (Bitan et al., 2003; Golde et al., 2000; Kuperstein et al., 2010; Scheuner et al., 1996). In the non-amyloidogenic pathway, APP is cleaved by α-secretase producing sAPPα and αCTF (or C83).

Early work has revealed Aβ as the main constituent of senile plaques establishing its central role in AD pathophysiology (Cras et al., 1991; Eanes and Glenner, 1968; Glenner and Wong, 1984a, 1984b; Kang et al., 1987; Masters et al., 1985). Additionally, genetics and genomic studies have so far identified 52 pathogenic APP mutations including the Swedish (K670N/M671L), Florida (I716V), and London (V717I) mutations, all of which are located near the β-secretase or γ-secretase cleavage sites and are associated with increased Aβ accumulation in familial or early-onset AD (for review (Tcw and Goate, 2017). In addition, the Icelandic A673T mutation has recently been identified in Icelandic and Scandinavian populations and carriers have a significantly lower risk of developing AD (Jonsson et al., 2012). The protective effect of the A673T mutation is believed to be primarily achieved through decreased Aβ production (Martiskainen et al., 2017; Xia et al., 2021).

Over the past decades, several Aβ-based mouse models have been developed to study the role of Aβ in AD, such as mice carrying mutations in APP and PS1. The APP/PS1 model carries the Swedish APP mutation (K670N/M671L) and the PS1 mutation (M146V). The 5xFAD mice carry the Swedish (K670N/M671L), Florida (I716V), and London (V717I) mutations in the APP gene, as well as the M146L and L286V mutations in the presenilin-1 (PS1) gene and is one of the most frequently used and best characterised models of AD (Oakley et al., 2006). These mice develop robust amyloid plaque pathologies that are suggested to trigger synaptic and neuronal loss (Forner et al., 2021; Jawhar et al., 2012; Oakley et al., 2006; Oblak et al., 2021), inflammatory responses (Sil et al, 2021) and loss of synaptic proteins (Anschuetz et al., 2025b; Jang et al. 2025). The protective effect of the Icelandic A673T mutation has been studied *in vitro* (Jonsson et al., 2012; Kokawa et al., 2015; Maloney et al., 2014; Wittrahm et al., 2023), *in vivo* using Aβ injection models, as well as APP knock-in mice and rats (Célestine et al., 2024; Shimohama et al., 2024; Tambini et al., 2020). However, the effect of the Icelandic mutation in transgenic APP mice remains elusive. We therefore here performed histopathological, immunoblot and ELISA immunoassays to access whether the introduction of the Icelandic mutation in a 5xFAD background reduces Aβ levels and rescues subsequent Aβ pathologies *in vivo*.

## Materials and Methods

### Animals and study design

All animal experiments were performed in accordance with the European Communities Council Directive (63/2010/EU) with local ethical approval under the UK Animals (Scientific Procedures) Act (1986) and its amended regulations (2012), and under the project licence number PP2213334 compliant with the ARRIVE guidelines 2.0 (du Sert et al., 2020). The study was exploratory. No power calculations were performed a priori.

Mice were bred at the Medical Research Facility of the University of Aberdeen (Aberdeen, Scotland). Heterozygous 5xFAD mice, on a black C57Bl6/J background (B6.Cg Tg (APPSwFlLon, PSEN1*M146L*L286V; 6799Vas/Mmjax, JAX MMRRC Stock# 034848) were crossed with mice harbouring the Icelandic mutation generated by CRISP-Cas gene editing of a single nucleotide into the murine APP gene at position 673 on a black C57Bl6/J background, termed APP^A673T^ mice. Crosses were bred from heterozygous 5xFAD (male or female) with heterozygous APP^A673T^ (male or female). Ear biopsies were genotyped for the 5xFAD and the A673T mutation in the APP gene by Transnetyx Inc. (Cordova, USA) and yielded heterozygous offspring only. Mice were grouped by sex and according to one of the four genotypes: C57Bl6/J wild type (WT), APP^A673T^, 5xFAD and 5xFADx APP^A673T^. A total of seventy-one male and female mice, 6- to 7-month-old, were included in the study (Table 1). Experimental mice were kept in sex- and genotype-specific litters ≥2 in stock box open housing under constant environmental conditions (20-22°C temperature, 50-65% humidity, an air exchange rate of 17-20 changes per hour, and a 12-h light/dark cycle with lights turned on at 7 am with simulated sunrise/sunset) and *ad libitum* chow (Special Diet Services, Witham, UK) and water throughout. Mice were provided with corncob bedding, paper strips, and cardboard tubes (DBM Scotland Ltd, Scotland) as enrichment throughout the experiment. They were kept in the same holding room throughout the study except when they were transferred to the euthanasia room for sacrifice and tissue harvest. Experimenters and care takers were blinded to the genotype of mice during maintenance and tissue collection. Following tissue collection, independent experimenters, also blinded to the genotype of mice, performed immunohistochemistry, ELISAs, and all statistical analyses relating to these measurements.

**Table 1:**
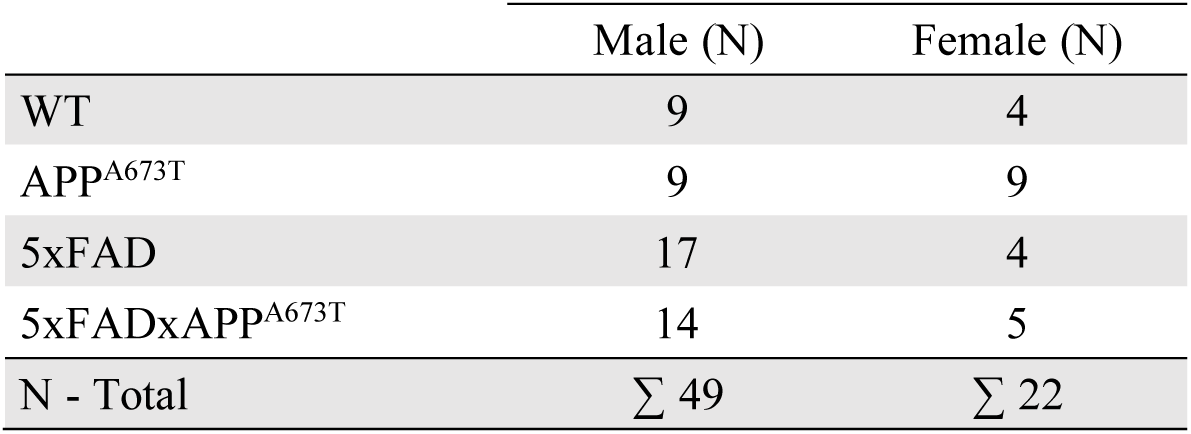
Study groups and cohort sizes. WT: C57Bl6/J wild-type mice, APP^A673T^: Icelandic mutation mice, 5xFAD: five familial Alzheimer’s disease mice, 5xFADxAPP^A673T^: crosses carrying both the 5xFAD mutations and the A673T mutation in the APP gene. N: number of mice. Mice were 6- to 7- month-old when they were perfused for tissue collection.

|  | Male (N) | Female (N) |
| --- | --- | --- |
| WT | 9 | 4 |
| APP<sup>A673T</sup> | 9 | 9 |
| 5xFAD | 17 | 4 |
| 5xFADxAPP<sup>A673T</sup> | 14 | 5 |
| N - Total | $\Sigma$ 49 | $\Sigma$ 22 |

### Animal perfusion and brain tissue collection

Brain tissue was harvested from all seventy-one mice (Table 1). All chemicals were purchased from Merck Millipore (Burlington, MA, USA) if not otherwise stated. Mice were euthanised via intraperitoneal injections of a lethal dose of sodium pentobarbital (Dolethal (200 mg/ml), Covetrus, UK) before undergoing intra-cardiac perfusion with heparinised phosphate-buffered saline (0.1 M PBS with 0.05% (v/w) heparin, pH 7.4) for 5 minutes. Skulls were dissected and whole brains retrieved. The right brain hemisphere was dissected, fixed overnight at room temperature in 10% (v/v) neutral-buffered formalin, dehydrated and embedded in paraffin. Sagittal sections were prepared at 5 µm using a rotary microtome (HM 325, Leica Biosystems, Sheffield, UK), and mounted onto glass slides (SuperFrost^TM^, Thermo Fisher Scientific, Lutterworth, UK). Sagittal sections were collected from regions at interaural 0.96 to 1.44 mm lateral of midline (Paxinos and Franklin, 2019), and three sections were collected on one slide for each mouse and antibody. After brain removal, the left-brain hemisphere was transferred immediately to liquid nitrogen and kept at −80°C until used for protein extraction, ELISA and immunoblot quantification.

### Aβ immunohistochemistry and quantification of Aβ plaques

Wax-embedded sagittal sections were stained in a sex-specific way using four immunohistochemistry staining boxes for male and two for female samples. Each box included a balanced number of all four genotypes. All chemicals were purchased from Merck Millipore (Burlington, MA, USA) unless otherwise stated. Sections were stained according to our standard protocol (Lemke et al., 2020) using the 6E10 anti-Aβ antibody (Biolegend # 803004, diluted 1:1,000). Images of hippocampal *cornu ammonis* (CA1), the dentate gyrus (DG), the visual cortex (CTX), the prefrontal cortex (PFC), and the cerebellum (CB) were taken using a light microscope at a 100x magnification (Axio Imager M1, Carl Zeiss, Jena, Germany) and saved as TIFF file format. Entire microscopic images were analysed using ilastik (Version 1.4.0.post1) (Berg et al., 2019) and Fiji (Version 2.14.0)(Schindelin et al., 2012). The pixel and object classification tool in ilastik enabled training of the software based on a small subset of samples and then apply them to larger sets of images) (Berg et al., 2019). Models were trained to segment images into positively stained pixels and unstained background tissue or artefacts, and additionally to specifically recognise extracellular Aβ plaques. After applying these models to all images, the percentage of positively stained area for the entire image, as well as extracellular plaques characteristics (number, size, and area) were quantified using Fiji. The total stained area (%),plaque count, average plaque size (µm^2^) and plaque area (µm^2^) were each analysed.

### NeuN immunohistochemistry

Wax-embedded sagittal sections were dewaxed and stained as described above using NeuN (Millipore #mAB377 diluted 1:1,000). Images from CA1, DG, CTX, PFC and CB were taken, and positive area was quantified as described above (percentage of positively stained area).

### Protein extraction

All chemicals were purchased from Merck Millipore (Burlington, MA, USA) unless otherwise stated. The left hemibrains were pulverized in a liquid nitrogen prechilled stainless steel mortar and pestle (BioPulverizer, BioSpec, Oklahoma, USA) and homogenized with a pestle and hammer. RIPA lysis buffer (Thermo Fisher Scientific, #89900) containing Pierce Protease and Phosphatase Inhibitor Mini Tablets (Thermo Fisher Scientific, # A32959) and 1mM AEBSF (4-(2-aminoethyl)benzenesulfonyl fluoride hydrochloride (Thermo Fisher Scientific #78431) were added in a ratio of 5:1 (mL buffer to mg wet tissue) and the homogenate was incubated for 30 minutes on ice with occasional agitation. After centrifugation at 19,000 g for 2 hours at 4 °C (Centrifuge 5427 R – Microcentrifuge, using the FA-45-48-11 rotor), the supernatant (referred to as the RIPA-soluble supernatant fraction S1) was transferred into new reaction tubes. The residual pellet was homogenised in 5 volumes TBS (pH 7.6) containing 5 M guanidine hydrochloride (GuHCl) and 1mM AEBSF and incubated with mild agitation (11 rotations per minute, Multi Bio RS-24, Biosan, Riga, Latvia) for 16 hours at room temperature. After centrifugation at 15,000 g for 30 minutes at room temperature, the resultant supernatant (referred to as GuHCl fraction or RIPA-insoluble fraction or S2) was transferred into new tubes. AEBSF was added to both S1 and S2 extraction buffers at a 1mM final concentration to prevent degradation of Aβ. S1 and S2 fractions were stored at −20 °C until use. Total protein concentration of S1 and S2 fractions was determined using the bicinchoninic acid (BCA) protein assay (Pierce™ BCA Protein Assay Kit, Thermo Fisher Scientific, #23225) with bovine serum albumin (BSA: 0.125 – 2.000 mg/ml) as a reference standard.

### Aβ, tau and synaptic proteins ELISA

All ELISAs were conducted according to the manufacturer’s instructions, and each sample was measured in duplicates.

RIPA-soluble S1 was used to measure human Aβ40 (Invitrogen #KHB3481), human Aβ42 (Invitrogen # KHB3441), mouse tau (Invitrogen #KMB7011), mouse synaptosomal associated protein 25kDa (SNAP25, MyBiosource #MBS451917), mouse syntaxin 1A (STX1A, MyBiosource #MBS452386), and mouse synaptophysin (SYP, MyBiosource #MBS453910). First, all S1 samples were diluted to a protein concentration of 4µg/µl in RIPA (including protease and phosphatase inhibitors + AEBSF). For Aβ40 and Aβ42, S1 samples were further diluted 1:5 in dilution buffer provided within each kit. All 5xFAD and 5xFADxAPP^A673T^ samples were used, and one WT and one APP^A673T^ sample was included on each plate as a control. For tau, S1 samples at 4µg/µl in RIPA were used, and quantification was conducted for all 71 mice. For synaptic proteins, S1 samples were further diluted in PBS at 1:2 for STX1A and 1:10 for SYP and SNAP25 and quantification was conducted for 70 mice (1 female WT excluded for SYP/SNAP25 due to sample preparation error). Additionally, Aβ40 and Aβ42 were quantified in GuHCl S2 fractions using the same kits as above. All S2 samples were first diluted to a protein concentration of 1µg/µl in TBS (pH 7.6) containing 5M GuHCl (including protease and phosphatase inhibitors + AEBSF) and further diluted 1:1,000 for Aβ40 or 1:7,500 for Aβ42 using the dilution buffer provided within each kit. All 5xFAD and 5xFADxAPP^A673T^ samples were used, and one WT and one APP^A673T^ sample was included on each plate as a control.

### Quantification of APP and APP fragments by immunoblotting

S1 RIPA-soluble samples were used for immunoblotting (4µg/µl in RIPA buffer including protease and phosphatase inhibitors + AEBSF). All chemicals were purchased from Merck Millipore (Burlington, MA, USA) if not otherwise stated. In brief, protein extracts were mixed with 4x Laemmli sample buffer (Bio-Rad Laboratories, CA, USA, #1610747) and incubated for 15 minutes at 37°C. Twenty µg protein per lane was loaded onto stain-free 4-15% gradient glycine gels (Bio-Rad Laboratories #4568086) and a protein standard (Bio-Rad Laboratories # 1610376) was loaded onto each gel as molecular weight (MW) marker. Proteins were separated in Tris-glycine-buffer (192 mM glycine, 25 mM Tris and 0.9% (w/v) SDS) at 100V for around 2 hours on ice using a Mini-PROTEAN Electrophoresis Cell (Bio-Rad Laboratories). Proteins were transferred to methanol-activated PVDF membranes (Bio-Rad Laboratories #1620177) at 5V for 30 minutes in Towbin transfer buffer (25 mM Tris, 200 mM glycine, 0.1% (w/v) SDS and 20% (v/v) ethanol. Membranes were then blocked for 1 h at RT in blocking solution (4% (w/v) BSA in TBS-T (TBS with 0.2% (v/v) Tween-20) and incubated overnight at 4°C in 5 mL primary antibody (Table 2) diluted in blocking solution. The next day, membranes were washed 3 × 10 minutes in TBS-T and incubated for 1 h at RT in 25mL secondary antibody (goat anti-mouse IgG, Bio-Rad Laboratories #5178-2504, or goat anti-rabbit IgG, Bio-Rad Laboratories #5196-2504, 1:5,000) diluted in blocking solution containing StrepTactin-HRP conjugate (Bio-Rad Laboratories #1610381; 1 µL conjugate per 100 mL blocking solution). After washing 3 × 10 minutes in TBS-T, membranes were overlaid for 1 min with ECL solution (Bio-Rad Laboratories # 1705061). The chemiluminescence signals were detected by the ChemiDoc Imaging System and the Image Lab software and normalised to protein loading signals using Coomassie Blue stain (0.1% Coomassie in 20% acetic acid and H_2_O). A mixture of all samples was included on each gel for between-gel normalization.

**Table 2:**
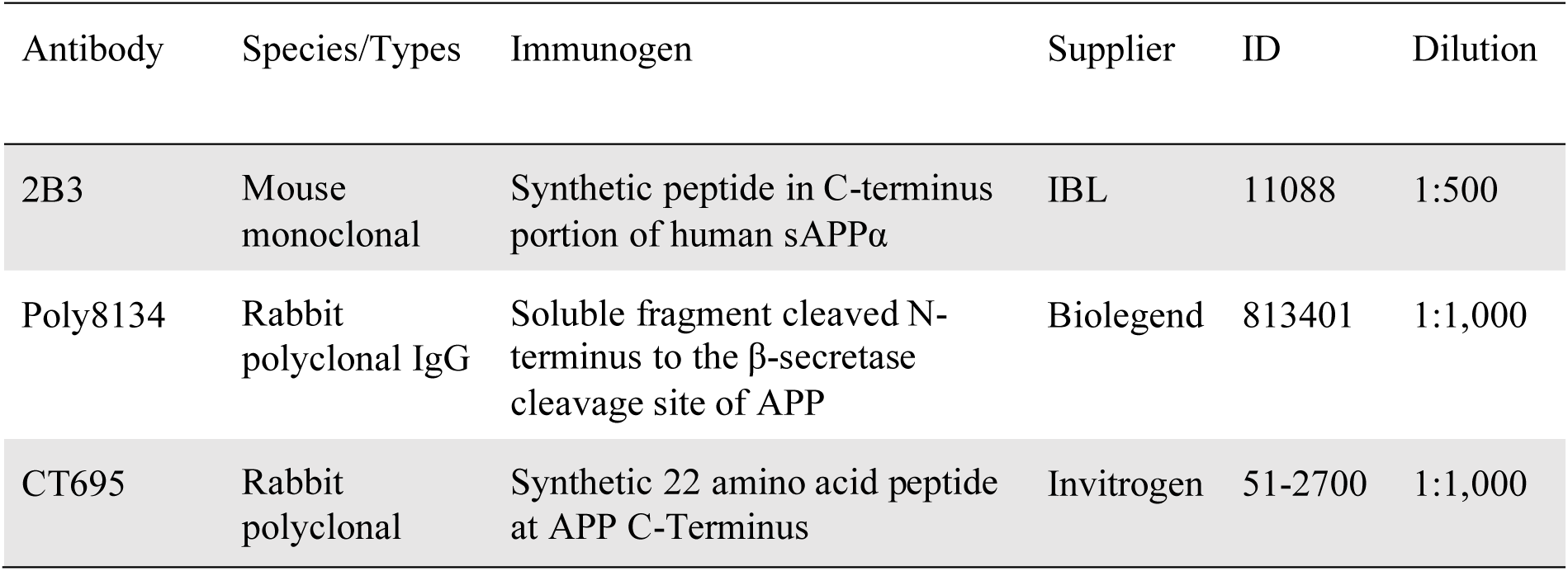
List of antibodies. Primary anti-Aβ antibodies used for immunoblotting.

| Antibody | Species/Types | Immunogen | Supplier | ID | Dilution |
| --- | --- | --- | --- | --- | --- |
| 2B3 | Mouse monoclonal | Synthetic peptide in C-terminus portion of human sAPPα | IBL | 11088 | 1:500 |
| Poly8134 | Rabbit polyclonal IgG | Soluble fragment cleaved N-terminus to the β-secretase cleavage site of APP | Biolegend | 813401 | 1:1,000 |
| CT695 | Rabbit polyclonal | Synthetic 22 amino acid peptide at APP C-Terminus | Invitrogen | 51-2700 | 1:1,000 |

### Data analysis

No a priori exclusion criteria were set. However, some IHC samples were excluded due to tissue damage during sectioning or lack of staining possibly due to sample preparation errors, and additionally some samples were excluded after immunoblotting due to damage of the gel. Details are specified in the respective sections below. Data were analysed and graphs generated in R (Version 4.4.3, R Core Team, Vienna, Austria) using linear models or generalized linear models and analysed using 2- or 3-Way ANOVA or Wald χ² tests. Where appropriate, post-hoc tests were performed using Bonferroni correction. For 6E10 and NeuN staining, males and females were analysed separately and the effects of brain region, genotype and their interaction on each parameter were assessed. For each analysis, it was first determined whether data met assumptions for normality or if any data transformations were necessary. Data met necessary assumptions after transformation using either simple methods (square root, log) or more advanced methods (Box-Cox or Yeo-Johnson transformation). As IHC was performed over several days, a nuisance factor “Staining Day” was included in statistical models if it had a significant effect on the variable being analysed. Total 6E10-positive area showed a significant nuisance factor effect in both males and females. Meanwhile in the analysis of plaque parameters and NeuN positive area, the staining day showed only a weak or non-significant effect and was therefore excluded as a factor.

A similar approach was taken for ELISA data, where the effects of sex, genotype and their interaction on protein levels were assessed. Data were first tested for necessary assumptions and transformed if necessary. For Aβ and tau ELISA. data were transformed using either simple or more advanced methods (see above) while synaptic protein data already met assumptions for two-way ANOVA. Due to the large number of samples multiple ELISAs were performed, which in part were from different lots and performed on different days. This was accounted for by inclusion of a nuisance factor where necessary. Nuisance factor was included in S1 Aβ42 and analysis of synaptic proteins.

Western blot data were analysed using one-way ANOVA with factor genotype following data transformation where necessary (for details, see figure legends). All statistical outcomes are reported based on linear or generalised linear models of transformed data, but figures show untransformed data. Due to a sample preparation error 1 sample (female WT) had to be excluded from SYP and SNAP25 ELISA. No other samples or data points were excluded from analysis. For each genotype and sex Pearson correlation matrices were generated from Aβ ELISA and Aβ IHC data and compared visually and statistically using the Jennrich test (Jennrich, 1970) to determine if the matrices were significantly different from each other. To determine whether the level of soluble or insoluble Aβ42/Aβ40 affected plaque counts and whether this effect varies between genotypes, generalized linear modelling was used. Negative binominal models were used and nested models (with or without interaction/factors) were compared using likelihood ratio tests to determine significance of each main effect (Aβ42/Aβ40 ratio and genotype) and interaction. Similarly, linear modelling was applied to determine the effect of Aβ42/Aβ40 ratio on plaque area and whether this differs between genotypes. All data are presented as mean ± standard deviation (S.D.) and alpha was set to p < 0.05.

## Results

We have experienced increased mortality in female 5xFAD mice during cohort aging (data not shown). The survival rate (until tissue harvest) was lowest in female 5xFAD (57%) compared to all other genotypes/sexes (between 80 and 100%). The remaining experimentally used mice were generally in good health when they were investigated at the age of 6 months (normal activity, no piloerection etc.). Furthermore, body weights differed considerably between genotypes (F_Genotype_(3,63) = 4.86, *p* = 0.0042) and sexes (F_sex_(1,63) = 133, *p* < 0.0001). In male cohorts, 5xFAD and 5xFADxAPP^A673T^ were generally lighter than WT and APP^A673T^ mice (WT: 35.23±2.59 g; APP^A673T^: 35.80±3.76 g; 5xFAD: 32.47±3.43 g; 5xFADxAPP^A673T^ 33.79±3.83 g). This was also the case in female cohorts (WT: 25.98±0.82 g; APP^A673T^: 26.93±2.55 g; 5xFAD: 22.60±1.45 g; 5xFADxAPP^A673T^ 23.20±0.84 g).

### Icelandic mutation and Aβ pathology

We proceeded to assess via IHC whether the introduction of the Icelandic mutation in a 5xFAD background changed Aβ levels using the monoclonal antibody 6E10. This antibody is widely used in AD research; it recognises APP fragments that contain the Aβ sequence (including full-length Aβ40 and Aβ42, as well as smaller fragments when used during immunoblotting) and is expected to label both intracellular and extracellular deposits of APP and Aβ. Representative micrographs of male 5xFAD and 5xFADxAPP^A673T^ (Fig. 1A), female 5xFAD and 5xFADxAPP^A673T^ (Fig. 1B), as well as WT and APP^A673T^ mice (Supplementary Fig. S1A) reveal uniform and punctate cytosolic staining (Fig. 1A, B & Fig. S1A, black arrowheads) and frequent nuclear as well as occasional axonal/ dendritic staining (Fig. 1A, B & Fig. S1A, white arrowheads). In WT and APP^A673T^ mice (Fig. S1A), there were abundant 6E10-positive neurones across all cortical layers in visual cortex and PFC and especially in the pyramidal cell layer of CA1 and granular cell layer of DG. Fewer 6E10-positive cells were found in other CA1 and DG layers as well as in the hilus. In cerebellum granule cell layer showed widespread cytoplasmic labelling while fewer positive cells were seen in the molecular layer. Additionally, large Purkinje cells were also frequently positive for 6E10 labelling. A similar cytosolic and axonal/ dendritic staining was also seen in 5xFAD and 5xFADxAPP^A673T^ (Fig. 1A, B).

**Figure 1:**
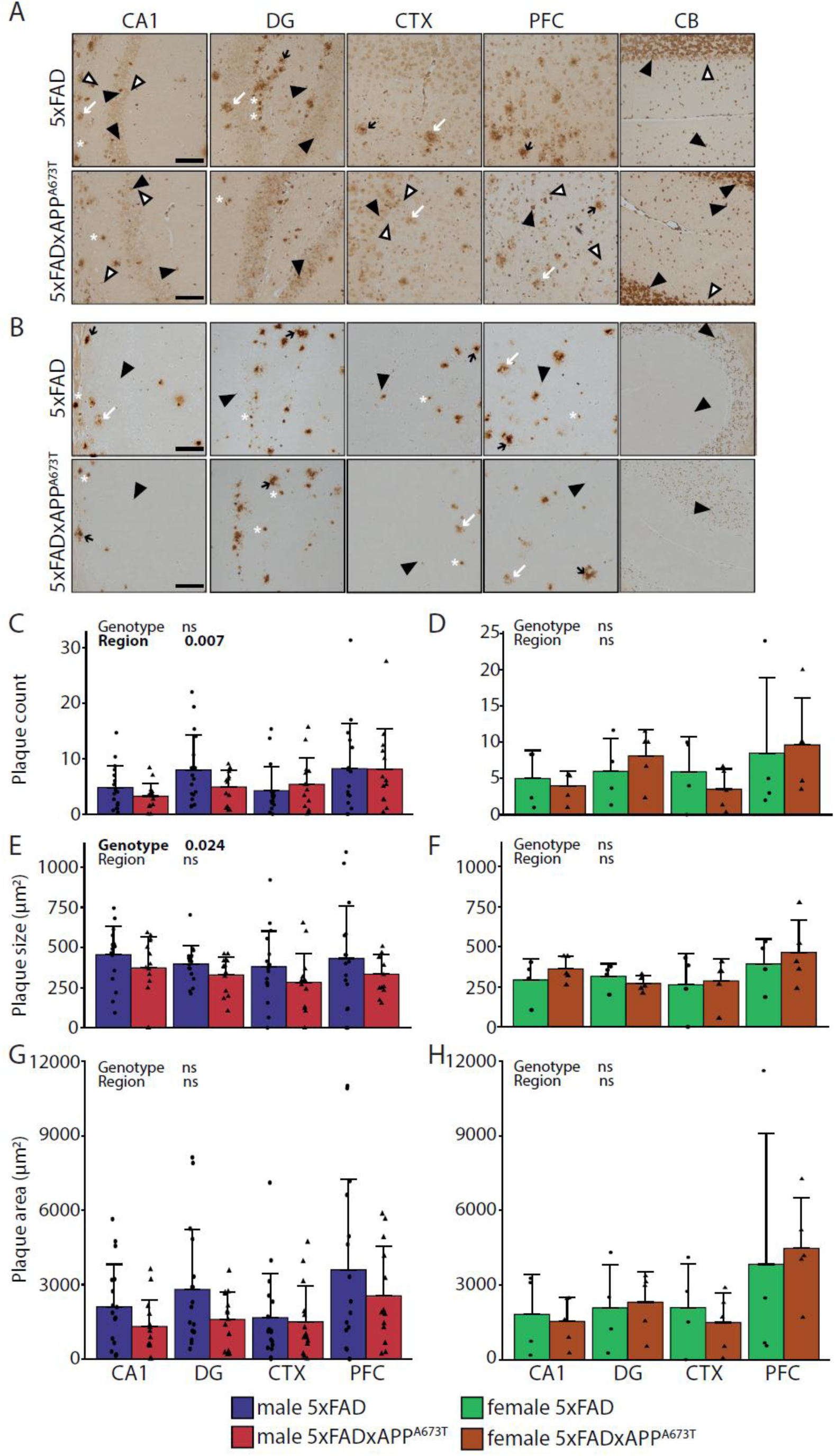
Aβ immunohistochemistry using the antibody 6E10. (A) Representative Aβ immunohistochemistry images of brain sections of male 5xFAD and 5xFADxAPP^A673T^ mice stained with the antibody 6E10 (Biolegend # 803004, diluted 1:1,000). Images from CA1, DG, CTX and PFC were taken using a light microscope at a 100x magnification. *Black arrowheads, cytosolic staining; white arrowheads, axonal/dendritic staining; black arrows, dense core plaques with halo; white arrows, small dense plaques with no/ little halo; asterisk, diffuse plaques; scale bars, 100µm*. 6E10 labelling was quantified using ilastik for plaque counts (C&D), plaque size (E&F) and plaque area (G&H) in male and female 5xFAD and 5xFADxAPP^A673T^ mice in four brain regions. Data is shown as individual values, group mean, and S.D. Statistical analysis entailed Wald χ² test (B&C) or two-way ANOVA (E-H) with genotype and region as independent variables. No data transformation was performed for plaque counts (C,D) while size and area were Box-Cox (E,F,H) or square-root transformed (G). Males: 5xFAD: n=17 (PFC n=16), 5xFADxAPP^A673T^: n=14 (PFC n=13). Females: 5xFAD: n=4, 5xFADxAPP^A673T^: n=5. Abbreviations: CA1: hippocampal CA1, CTX: visual cortex, DG: dentate gyrus, ns: not significant, PFC: prefrontal cortex.

Extracellular Aβ deposits were absent in WT and APP^A673T^ mice (Fig. S1A), but 5xFAD and 5xFADxAPP^A673T^ mice of both sexes showed abundant extracellular Aβ deposits (Fig. 1A, B). These consisted of characteristic Aβ plaques with an intensely labelled core and a fainter diffuse halo (Fig. 1A, B, black arrows). In addition, deposits of smaller, intensely labelled core-only plaques with little to no halo (Fig. 1A, B, asterisk) and less intensely labelled diffuse plaques with no discernible core (Fig. 1A, B & Fig. S1A, white arrows) were identified. All three types of plaques were found in hippocampal and cortical areas in 5xFAD and 5xFADx APP^A673T^, but none were seen in CB (Fig. 1A, B). Plaque number, size and area were measured to quantify extracellular Aβ deposits. These three parameters differed significantly between the four genotypes, confirming the Aβ plaque pathology phenotype in 5xFAD and 5xFADxAPP^A673T^ male and female crosses (Fig. S1B-G, *p* values < 0.001). When the total 6E10 signal was quantified, these genotype differences persisted only in female but not male cohorts (Fig. S1H, *p* not significant in males and Fig. S1I, *p* < 0.001 in females). However, while the number of plaques was similar between 5xFAD and 5xFADxAPP^A673T^ male (Fig. 1C) and female mice (Fig. 1D), male 5xFADxAPP^A673T^ had significantly smaller plaques than male 5xFAD (Fig. 1E, F_Genotype_(1,114) = 5.24, *p* = 0.024), but no genotype-related differences were measured for this parameter in female cohorts (Fig. 1F). The plaque area was also similar between genotypes in male (Fig. 1G) and female mice (Fig. 1H). Finally, the number of plaques varied significantly between brain regions in male 5xFAD and 5xFADxAPP^A673T^ males (Fig. 1C, χ²_Brain Region_(3) = 12.14, *p* = 0.007), where significantly more plaques were counted in PFC than in CA1 (*post-hoc* test *p* = 0.009).

In summary, we have confirmed the Aβ plaque pathology phenotype in 5xFAD and 5xFADxAPP^A673T^ male and female cohorts and show for males that the A673T mutation significantly decreases the size of Aβ plaques in 5xFADxAPP^A673T^ crosses compared to 5xFAD.

Given this significant reduction of Aβ plaque size in male 5xFADxAPP^A673T^ crosses, we further explored, using ELISA, whether this led to changes in soluble and insoluble Aβ40 and Aβ42 isoforms (Fig. 2). While all 5xFAD and 5xFADxAPPA673T samples were measured, only one WT and one APP-APP^A673T^ samples were included. Both presented with very low signals, or signals below detection thresholds and confirmed the specificity of the ELISA assays for human Aβ (data not shown). Female 5xFAD and 5xFADxAPP^A673T^ crosses had almost twice as much soluble Aβ40 than their male counterparts (Fig. 2A, F_Sex_(1,36) = 12.77, *p* = 0.001), and this was also the case for soluble Aβ42 (Fig. 2B, F_Sex_(1,35) = 8.35, *p* = 0.007), while the Aβ42/Aβ40 ratio was similar between cohorts (Fig. 2C). Similarly, females of both genotypes had more insoluble Aβ40 (Fig. 2D, F_Sex_(1,36) = 7.67, *p* = 0.008), and Aβ42 (Fig. 2E, F_Sex_(1,36) = 8.02, *p* = 0.007), but again a similar Aβ42/Aβ40 ratio (Fig. 2F) compared to their male counterparts. Neither soluble, nor insoluble Aβ40 and Aβ42 nor their ratios differed between 5xFAD and 5xFADxAPP^A673T^ crosses, but a trend towards reduction for Aβ42/Aβ40 in S2 was observed for 5xFADxAPP^A673T^ compared to 5xFAD (Fig. 2F, F_Genotype_(1,36) = 3.31, *p* = 0.077).

**Figure 2:**
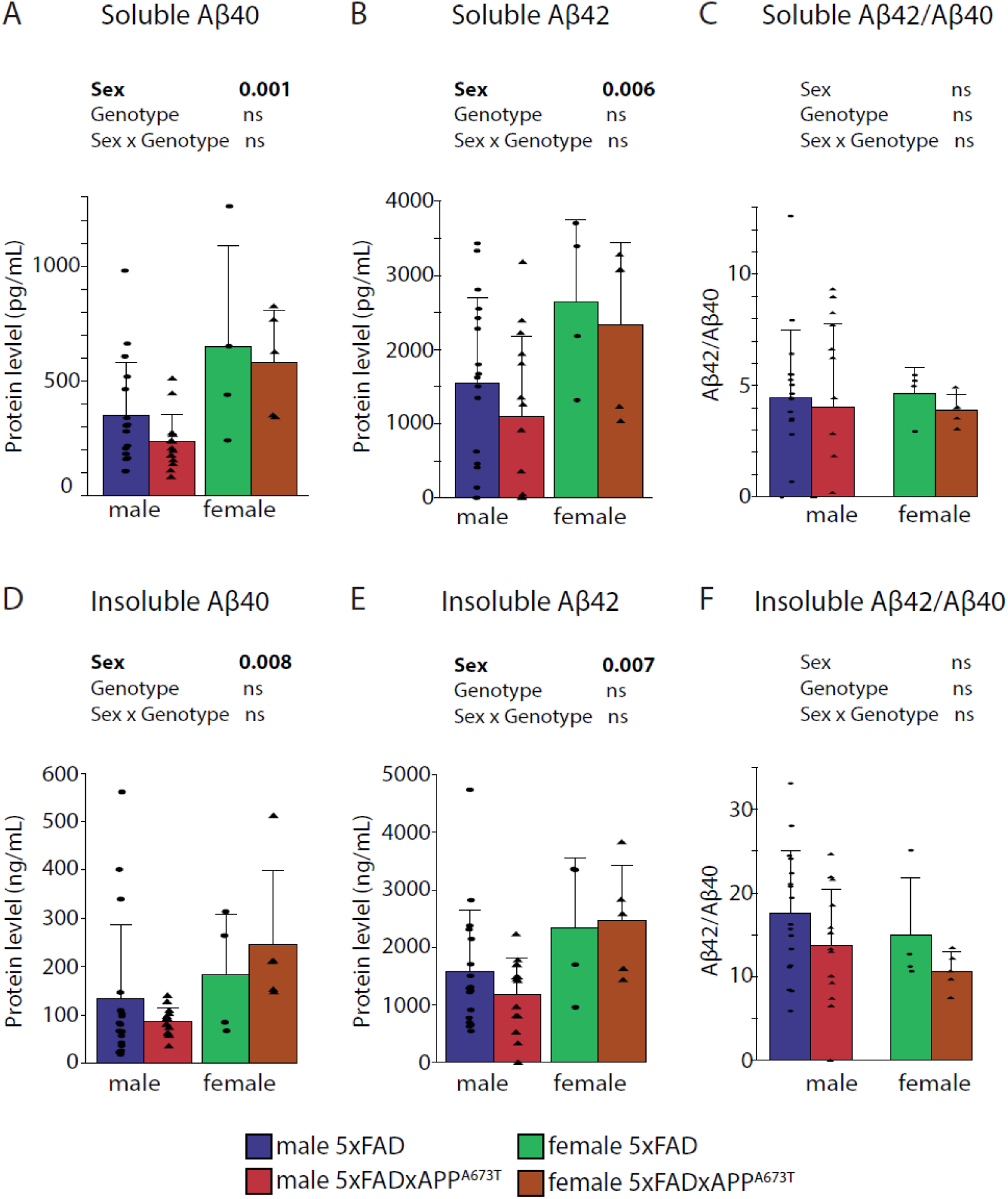
Quantification of soluble and insoluble Aβ. Human Aβ40 (Invitrogen #KHB3481) and Aβ42 (Invitrogen, #KMB3441) and Aβ42/Aβ40 ratios were quantified in RIPA-soluble (A-C) and insoluble fractions (D-E) in male and female WT, APP^A673T^, 5xFAD and 5xFADxAPP^A673T^ mice. Data were analysed using two-way ANOVA with sex and genotype as independent variables. Ratios did not require data transformation and remaining data were transformed using Yeo-Johnson (A, D) or Box-Cox (B, E) transformation. Data is shown as individual values, group mean, and S.D. Males: 5xFAD: n=17, 5xFADx APP^A673T^: n=14. Females: 5xFAD: n=4, 5xFADxAPP^A673T^: n=5. One wild-type and one APP^A673T^ were included on each ELISA plate as a control (see Methods). Abbreviations: ns: not significant.

We next investigated Aβ, APP, and its metabolites using immunoblotting to assess whether the A673T mutation would shift its processing from the amyloidogenic to the non-amyloidogenic pathway (Fig. 3, and additionally see Fig. S2 for uncropped images of the complete cohort). We have used three different anti-APP/Aβ antibodies (Table 2) on male cohorts as only these returned genotype-specific differences for Aβ plaques (Fig. 1).

**Figure 3:**
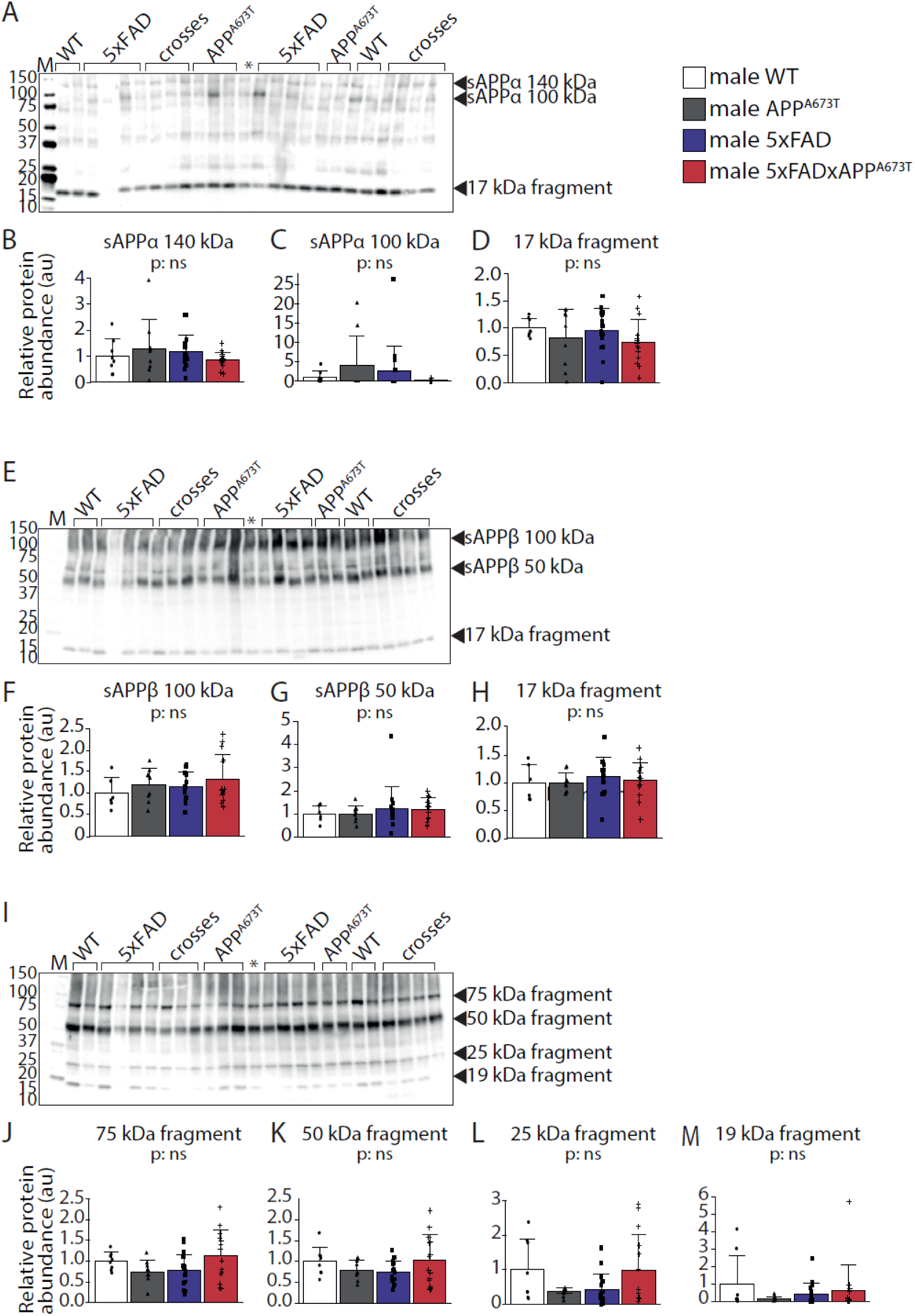
Quantification of APP/Aβ species using immunoblotting. Proteins from RIPA-soluble S1 fractions were separated by SDS-PAGE (20 µg per lane) and labelled with the antibody 2B3 against sAPPα (A), Poly8134 against sAPPβ (E), and CT695 against CTFs (I). Densitometric quantification of the bands of interest (arrowheads) was conducted using the Image Lab software and normalisation to total protein loading. For antibody 2B3, three bands at 10 kDa (B), 100 kDa (C) and 17 kDa (D) were identified. For the antibody Poly8134, three bands were quantified at 100 kDa (F), the 50 kDa (G) and 17 kDa (H). The third antibody, CT695 revealed four bands at 75 kDa (J), 50 kDa (K), 25 kDa (L) and 19 kDa (M). Data is shown as individual values, group mean, and S.D. Data were analysed using 1-way ANOVA with genotype as independent variable. No data transformation was required. Males: WT: n=8 (n=6 for Poly8134 antibody), APPA673T: n=9, 5xFAD: n=17 (n=15 for Poly8134 antibody), 5xFADx APP^A673T^: n=14. Abbreviations: ns: not significant.

The monoclonal antibody 2B3 (IBL # 11088) is directed against the C-terminus of human sAPPα. Applying our immunoblotting protocol to RIPA-soluble S1 fractions, this antibody revealed three bands: two higher molecular weight bands at around 140 and 100 kDa (sAPPα-140 and sAPPα-100), as well as a 17-kDa fragment (Fig. 3A, see black arrowheads). The levels of these three bands, however, was similar between genotypes (Fig. 3B-D). The second antibody, Poly8134 (Biolegend #813401), is polyclonal and directed against APPβ. It too revealed three bands: sAPPβ-100 (MW ∼100 kDa), sAPPβ-50 (MW ∼50 kDa) and a 17-kDa fragment (Fig. 3E, see black arrowheads), all of which were similar across the four genotypes (Fig. 3F-H). The third antibody, CT695 (Invitrogen #51-2700), reacts with CTFs of human APP and revealed four fragments: CTF75 (∼75 kDa), CTF50 (∼50 kDa), CTF25 (∼25 kDa), and CTF19 (19 kDa, Fig. 3I, see black arrowheads). Again, all these four bands were similar in quantity between genotypes (Fig. 3J-M). All three antibodies revealed considerable cross-reactivity for murine and human APP and metabolites, likely because mouse and human APP differ by only three amino acids (Serneels et al., 2020).

### Icelandic mutation, tau and synaptic proteins

Given the synergetic and reciprocal regulatory effect of Aβ and tau, and their established role in inducing synaptic protein alterations in AD patients and AD mouse models, we have further examined whether the A673T mutation in the APP gene changes endogenous tau levels and/or rescue alterations of synaptic proteins. Mouse tau and three synaptic proteins – SYY, SNAP25, and STX1A – were measured using mouse specific ELISAs (Fig. 4). Tau was similar across genotypes and sexes (Fig. 4A), as were SYP (Fig. 4B) and SNAP25 (Fig. 4C, all F values < 1). STX1A however, was different between the 4 genotypes (Fig. 4D, F_Genotype_(3,62) = 3.1, *p* = 0.034), but none of the differences reached statistical significance in *post-hoc* tests.

**Figure 4:**
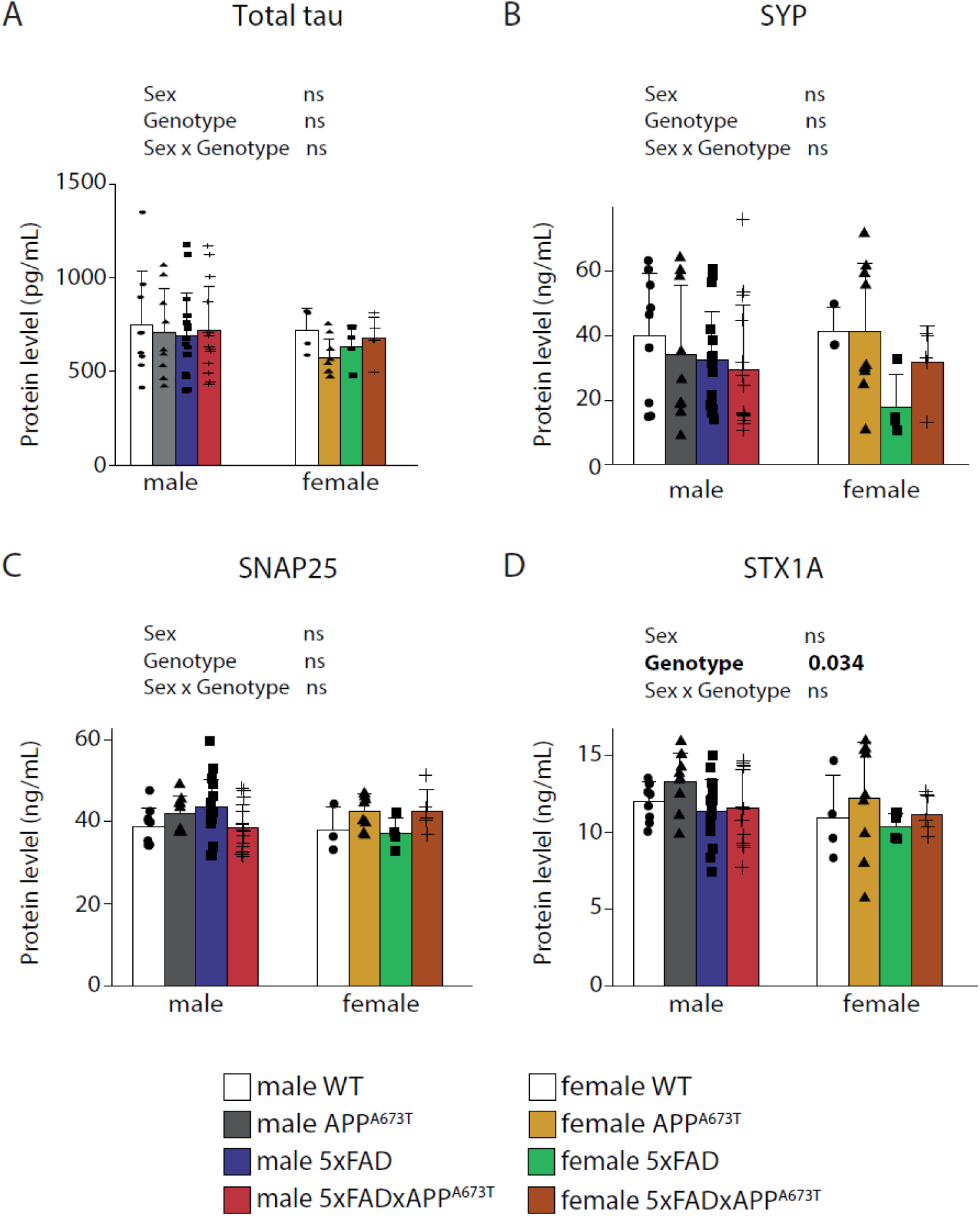
Quantification of tau and synaptic proteins. (A) Mouse tau (ELISA kit, Invitrogen #KMB7011), (B) mouse synaptophysin (SYP ELISA kit, MyBiosource #MBS453910), (C) mouse synaptosomal associated protein 25kDa (SNAP25, ELISA kit MyBiosource #MBS451917), and (D) mouse syntaxin 1A (STX1A, ELISA kit MyBiosource #MBS452386) were quantified in RIPA-soluble S1 fractions in male and female WT, APP^A673T^, 5xFAD and 5xFADxAPP^A673T^ mice. Data were analysed using two-way ANOVA with sex and genotype as independent variables. No data transformation was needed. Males: WT: n=9, APP^A673T^: n=9, 5xFAD: n=17, 5xFADx APP^A673T^: n=14. Females: WT: n=4 (SYP, SNAP25 and STX1A n=3), APP^A673T^: n=9, 5xFAD: n=4, 5xFADx APP^A673T^: n=5. Abbreviations: ns: not significant, SNAP25: synaptosomal associated protein 25kDa, STX1A: syntaxin 1A, SYP: synaptophysin.

### Icelandic mutation and prediction of amyloid pathology

Pearson correlations were generated for data from 5xFAD and 5xFADxAPP^A673T^ male and female mice. These correlations matrices included Aβ pathology (IHC and ELISA) and tau quantification (Fig. S3, see supporting information). Correlation matrices differed significantly between male 5xFAD and 5xFADxAPP^A673T^ mice (Fig. S3A and Fig. S3B, *p* < 0.001, see supporting information). Although differences were obvious between female 5xFAD and 5xFADxAPP^A673T^, sample sizes were too small to compare correlation matrices statistically (Fig. S3C and Fig. S3D, see supporting information). Overall, there was a high degree of correlation for Aβ (IHC with ELISA), especially in 5xFAD males, while almost no correlations were observed between Aβ and tau in either genotype. When only amyloid pathologies are correlated (Fig. 5A-D), we found that male 5xFAD mice showed significant positive correlations between Aβ40 and Aβ42 levels with plaque counts and plaque area which were almost entirely absent in 5xFADxAPP^A673T^ (Fig.5A and Fig.5B, see asterisks for significant correlations). Additionally, 5xFAD males showed significant negative correlations between Aβ42/Aβ40 ratio in S2 with plaque counts/area. In female mice, the Aβ42/Aβ40 ratio in S1 fraction correlated significantly with plaque count/area in 5xFAD mice, but this was not the case in 5xFADxAPP^A673T^ (Fig.5C and Fig.5D, see asterisks for significant correlations).

**Figure 5:**
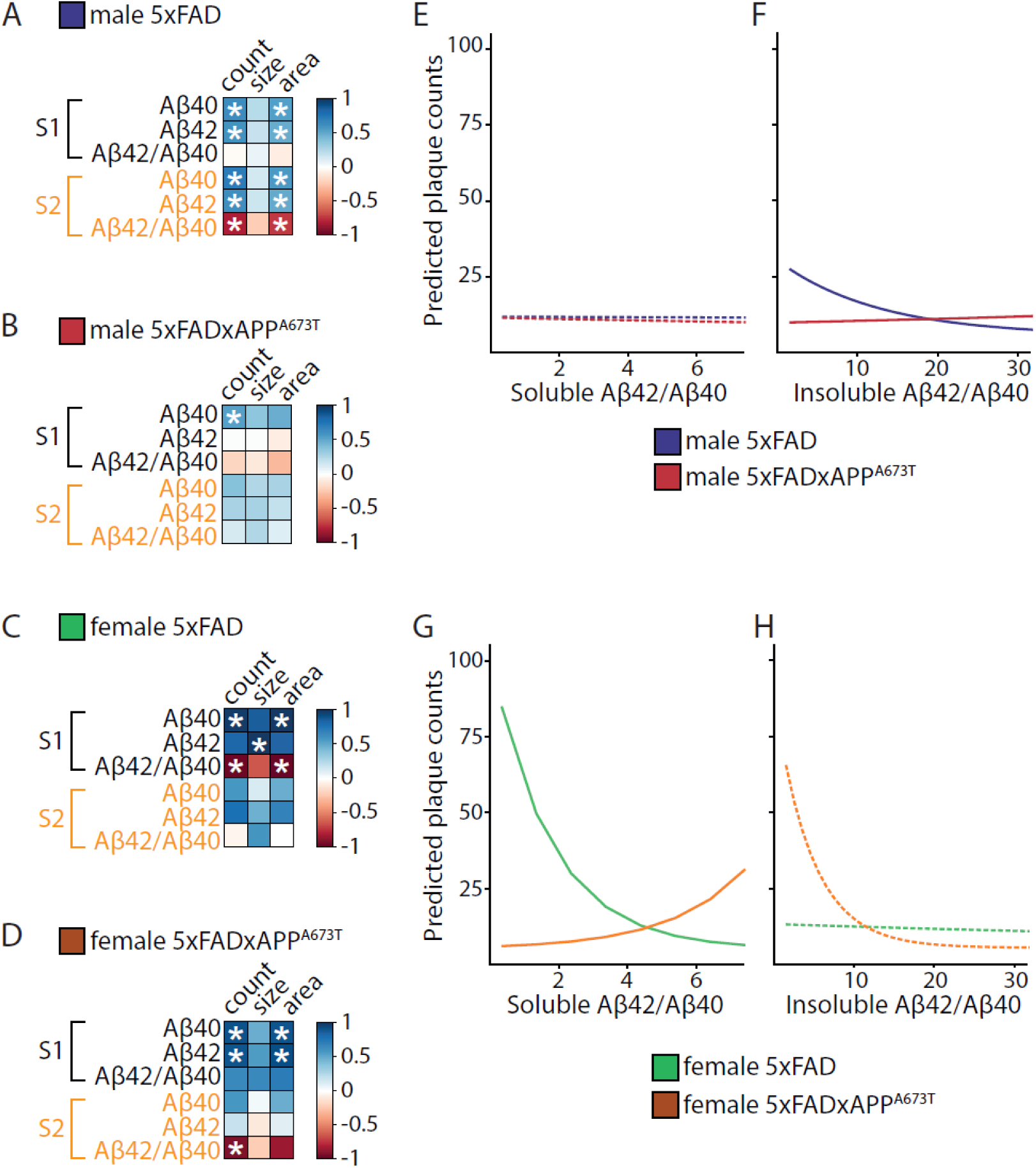
Correlation matrices and linear modelling for the different Aβ measurements. (A-D) Pearson correlation matrix between Aβ40 and Aβ42 levels and their ratio (Aβ42/ Aβ40) in soluble and insoluble fractions and plaque measurements (count, size, and area) are displayed for 5xFAD male (A), 5xFADxAPP^A673T^ male (B), 5xFAD female (C) and 5xFADxAPP^A673T^ female (D) mice with blue for positive correlations, red for negative correlations and white where no correlation was seen (* p < 0.05). Aβ levels were quantified using ELISA and plaque counts, size and total area were quantified using immunohistochemistry (averaged across brain regions). Data were analysed using Jennrich test to detect differences between matrices. (E-H) Generalized linear modelling to explore the effect of Aβ42/Aβ42 ratio in S1 and S2 on plaque counts and whether this differed across genotypes. Model-predicted plaque counts depending on Aβ42/Aβ40 ratio for 5xFAD and 5xFADxAPP^A673T^ male in S1 (E) and S2 (F), as well as 5xFAD and 5xFADxAPP^A673T^ female in S1 (G) and S2 (H) are presented. Dashed lines indicate non-significant effects.

To further explore these differences in correlations, generalised linear modelling was used to determine whether the Aβ42/Aβ40 ratio in insoluble and soluble fractions would predict plaque count and whether this effect differs between genotypes (Fig. 5E-H). In males, independent of genotype, the Aβ42/Aβ40 ratio in S1 did not influence plaque count (Fig. 5E). By contrast, in S2, 5xFAD males showed a negative association between Aβ42/Aβ40 ratio and plaque count (Fig.5F, *p* < 0.001). The relationship showed a positive direction in 5xFADxAPP^A673T^(p<0.001), resulting in lower plaque counts in 5xFADxAPP^A673T^ than 5xFAD males when the Aβ42/Aβ40 ratio is low, with a significant difference between both genotypes for the number of plaques which depended on Aβ42/Aβ40 (Fig.5F, likelihood ratio test: χ²(1) = 8.79, *p* = 0.003). In 5xFAD female mice, increase in Aβ42/Aβ40 ratio in S1 was associated with a predicted decrease in plaque counts (Fig.5G, *p* < 0.001). The opposite was the case in 5xFADxAPP^A673T^ females, with increasing Aβ42/Aβ40 values associated with an increase in plaque counts (Fig.5G, *p* =0.001). This resulted in lower predicted plaque counts in 5xFADxAPP^A673T^ compared to 5xFAD females for low values of Aβ42/Aβ40, and a significant difference between genotypes in the prediction of plaque count based on the Aβ42/Aβ40 ratio (Fig. 5G, likelihood ratio test: χ² (1) = 7.52, *p* = 0.0061). The Aβ42/Aβ40 ratio in S2 was not significantly associated with plaque counts in females independent of genotype (Fig.5H). The same patterns were seen when investigating the relationship between Aβ42/Aβ40 and total plaque area (Fig. S4).

### Icelandic mutation and neurodegeneration

Neuronal loss and gliosis associated with Aβ plaque pathologies have been reported for 5xFAD mice as early as 6 months of age (Jawhar et al., 2012; Oakley et al., 2006). Therefore, neurons were quantified in different regions of the brain using NeuN as a marker. This was done in 5xFAD, 5xFADxAPP^A673T^ crosses, as well as their control counterparts WT and APP^A673T^ (Fig. 6).

**Figure 6:**
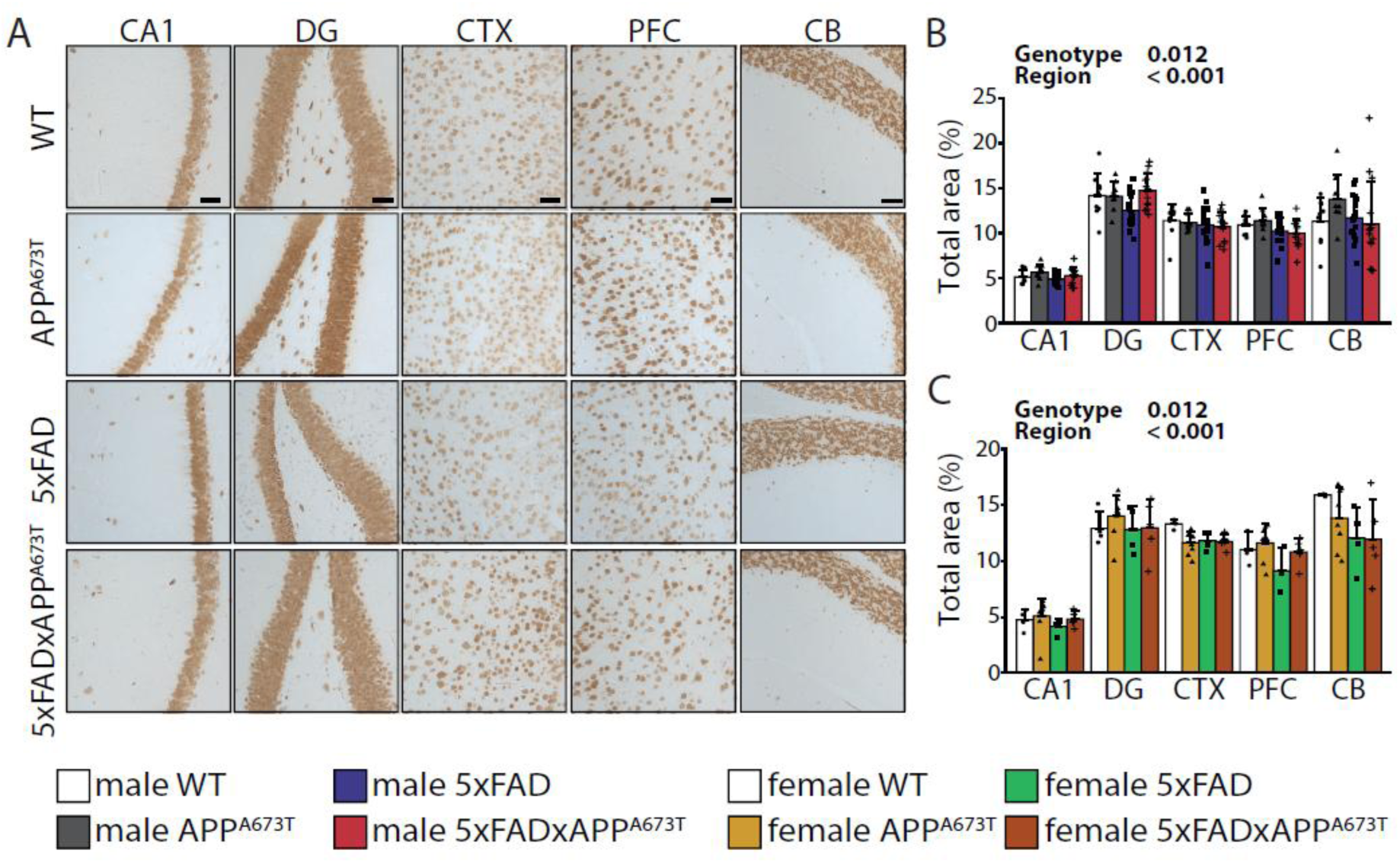
NeuN immunohistochemistry. (A) Representative NeuN immunohistochemistry images in brains of male WT, APP^A673T^, 5xFAD and 5xFADxAPP^A673T^ mice stained with the antibody NeuN (Millipore #mAB377 diluted 1:1,000). Images from CA1, DG, CTX and PFC were taken using a light microscope at a 100x magnification. *scale bars, 100µm*. NeuN levels were quantified using ilastik as total stained are in male (B) and female (C) mice in four individual brain regions. Data are shown as individual values, group mean, and S.D. Statistical analysis entailed two-way ANOVA with genotype and region as independent variables. Data were transformed using square-root transformation. Males: WT: n=9, A673T: n=9, 5xFAD: n=17 (PFC n=16), 5xFADxAPP^A673T^: n=14 (CTX and PFC n=13). Females: WT: n=4 (CB and PFC n=3), A673T: n=9 (PFC n=7, CB n=8), 5xFAD: n=4 (PFC n=3), 5xFADxAPP^A673T^: n=5. Abbreviations: CA1: hippocampal CA1, CTX: visual cortex, DG: dentate gyrus, ns: not significant, PFC: prefrontal cortex.

Representative NeuN images from CA1, DG, CTX, PFC, and CB are shown (Fig. 6A). Their quantification revealed significant genotype differences in male (F_Genotype_(3,222) = 3.72, *p* = 0.012) and female cohorts (F_Genotype_(3,84) = 3.00, *p* = 0.035). In males, the average over the five brain regions confirmed the difference between the genotypes (F_Genotype_(3,45) = 2.30, *p* = 0.090, data not shown) revealing a modest reduction of NeuN in 5xFAD compared to WT (−5.7%), while this reduction was even less pronounced in 5xFADxAPP^A673T^ crosses compared to WT (−2.8%). A similar, although not significant, observation was seen in females (data not shown), where NeuN was reduced in 5xFAD compared to WT (−12.3%), and again the reduction was less pronounced in 5xFADxAPP^A673T^ crosses compared to WT (−7.9%). Additionally, the NeuN signal differed significantly between brain regions both in male (Fig. 6B, F_Brain Region_(4,222) = 146.05, *p* < 0.001) and female cohorts (Fig. 6C, F_Brain Region_(4,84) = 90.75, *p* < 0.001). Post-hoc analyses yielded a lower NeuN signal in CA1 compared to all other regions in males (Fig. 6B, all *p* values < 0.001), and females (Fig. 6C, all *p* values < 0.001).

## Discussion

Here, we have investigated the effect of the protective Icelandic mutation, A673T, on Aβ pathology in the 5xFAD mouse model of AD (Oakley et al., 2006). FivexFAD mice were bred with APP^A673T^ mice resulting in 5xFADxAPP^A673T^ crosses, that are heterozygous for both the 5xFAD mutations and the A673T mutation in the APP gene. The overarching aim was to examine Aβ pathology, as well as tau and synaptic protein levels in 5xFAD and 5xFADxAPP^A673T^ mice, including their respective WT and APP^A673T^ controls. The main findings that we report are:

i. The A673T mutation significantly decreases the size of Aβ plaques in 5xFADxAPP^A673T^ male crosses compared to 5xFAD mice.
ii. Aβ40, Aβ42 and Aβ42/Aβ40 ratios were similar between 5xFAD and 5xFADxAPP^A673T^ crosses. However, the Icelandic mutation changed the association between Aβ42/Aβ40 plaque count/area: at low ratios, 5xFADxAPP^A673T^ tended to show lower predicted plaque burden than 5xFAD while the opposite was the case at high ratios.
iii. No differences were measured between 5xFAD and 5xFADxAPP^A673T^ crosses for Aβ immunoblot species, tau, synaptic proteins (SYP, SNAP25, and STX1A) or neuronal loss.

The pathological accumulation of Aβ, either caused by its decreased clearance and/or increased oligomerisation and aggregation, leads to synaptic alterations, neuroinflammation, and eventually neuronal cell death (Hampel et al., 2021). Several aggregation-promoting mutations have been identified near the β-secretase or γ-secretase cleavage sites in the APP gene (amyloidogenic APP pathway), such as the Swedish K670N/M671L, Florida I716V, or London V717I mutations. A mutation with opposite effects, the Icelandic A673T mutation, has been identified in Nordic populations, and carriers of this mutation have a significantly lower risk of developing AD presumably due to increased α-secretase cleavage (Jonsson et al., 2012; Martiskainen et al., 2017; Xia et al., 2021). In cellular models, A673T reduced amyloidogenic processing of APP and decreased Aβ aggregation by reducing the release of sAPPβ (Kokawa et al., 2015; Maloney et al., 2014). When the A673T was expressed in cell culture models expressing APP with the Swedish and London mutations, it reduced sAPPβ but Aβ42, Aβ40 and the Aβ42/Aβ40 ratio remained unchanged (Wittrahm et al., 2023) and it has also been shown in cells combining 29 FAD mutations with the A673T mutation that the protective effect of the A673T mutation was specific to certain mutations, e.g. the London mutation (V717I) but was absent in the Florida (I716V) and Swedish (KM670/671NL) mutations (Guyon et al., 2020). It was therefore reasonable to hypothesise that the Icelandic mutation A673T could counteract, at least in part, some of the effects introduced by the Swedish/Florida/London mutations in terms of Aβ and other subsequent pathologies in 5xFAD mice.

### Icelandic mutation and Aβ

Histopathologically, the A673T mutation led to a decrease in Aβ plaque size in 5xFADxAPP^A673T^ males compared to 5xFAD. While both Aβ40 and Aβ42 are found in plaques, however an increased cerebral Aβ42/Aβ40 ratio is another well-established biomarker of Aβ pathology in patients and 5xFAD mice, due to the greater aggregation propensity of Aβ42 (Andersson et al., 2025; Blennow and Zetterberg, 2018). While no overt differences were identified for soluble/insoluble Aβ40 and Aβ42, we found the way in which their ratio was associated with plaques differed considerably between 5xFAD and 5xFADxAPP^A673T^: male 5xFAD mice showed significant positive correlations between insoluble Aβ40 and Aβ42 with plaque counts & area. These were almost entirely absent in 5xFADxAPP^A673T^. Similarly, in females with heightened soluble and insoluble Aβ40 and Aβ42 levels (Sil et al. 2022, this study), the Aβ42/Aβ40 ratio in soluble fractions correlated significantly with plaque count/area in 5xFAD mice, but this was not the case in 5xFADxAPP^A673T^. When modelling these, genotype-differences depended on Aβ42/Aβ40 ratios; a protective effect (i.e. reduced plaque burden in 5xFADxAPP^A673T^) was seen at low ratios that disappeared or is reversed at high ratios. These differences suggest a genotype-dependent sensitivity to Aβ accumulation. They would also suggest the strength of the amyloid burden in 5xFAD mice is too aggressive and the protection is too weak to counteract their aggregation propensity.

Only a few publications have addressed the potential protective effects of the Icelandic mutation in AD models *in vivo*. The first used a knock-in rat model of humanized A673T-APP, K670N/M671L-APP (Swedish mutation) or both, and found a reduction of Aβ40 and Aβ42 pathology (using ELISA) for A673T-APP compared to wild-type APP but not when the Icelandic mutation was combined with the Swedish mutation (Tambini et al., 2020). Using immunoblotting they corroborated an increase in non-amyloidogenic APP metabolites (sAPPα) and a decrease in amyloidogenic APP metabolites (sAPPβ and βCTF) again for the Icelandic mutation alone, but not when combined with the Swedish mutation. The authors suggested that the Swedish and Icelandic mutations may act independently but the magnitude of the protective effect caused by the Icelandic mutation is smaller than the aggressive pathogenic effect of the Swedish mutation. We have found no differences in APP fragments between genotypes using immunoblotting, confirming a lack of efficacy of A673T when combined with the Swedish mutation and suggesting no shift in APP processing in 5xFAD mice when the A673T mutation is introduced on a Swedish/Florida/London background.

The second study generated knock-in mice with humanized APP with the Arctic (E693B) and Beyreuther/Iberian (I715F) mutations and compared them to mice also carrying the Icelandic mutation (Shimohama et al., 2024). The protective A673T mutation reduced plaque area in cortex and hippocampus at 8 months of age but at 12 months, only the number of plaques larger than 20 µm was decreased while smaller plaques showed similar levels in both genotypes. They additionally report a decrease in βCTF at 3 months (where no Aβ pathology is established yet) but it is unclear if this persists at older age, where we also could not see any shift in APP processing. In our male 5xFADxAPP^A673T^ mice, only the plaque size was decreased compared to 5xFAD, suggesting a more aggressive Aβ pathology produced by the Swedish/Florida/London mutations as compared to the Arctic or Beyreuther/Iberian APP mutations. This is also supported by *in vitro* findings, where it has been shown that the protective effect of the A673T mutation was specific to the London mutation (V717I) but was absent in the Florida (I716V) and Swedish (KM670/671NL) mutations (Guyon et al., 2020; Wittrahm et al., 2023).

Another study inoculated APPswe/PS1dE9 transgenic mice with either recombinant non-mutant human Aβ or human Aβ containing the A673T mutation once at 2 months of age and found no changes in Aβ levels when analysed at 6 months. There was only a rescue in synapse density and spatial memory which remained unexplained (Célestine et al., 2024). In this model, similar to our 5xFAD mouse, the role of PS1 mutations remain unexplored and individual contributions of these mutations to the amyloid load, and a possible block of the A673T protection are elusive to date. Lastly, a recent study that introduced the A673T mutation into a tau-transgenic model, L66, reported no modulation of Aβ and tau pathologies and no rescue of motor and neuropsychiatric behaviour in these mice (Anschuetz et al., 2025a).

5xFAD mice overexpress randomly integrated mutant human Aβ, while in APP^A673T^ mice, the Icelandic A673T mutation was generated in the murine APP gene. There is no clear evidence that murine A673T APP could affect human APP processing. However, It has been shown that co-expression of murine APP can alter Aβ pathology in APP23 transgenic mice but not in the much faster Aβ-depositing APPPS1 transgenic mice (Mahler et al., 2015). Moreover, the targeted knock-in of human BACE1 lead to amyloidosis purely based on murine Aβ (Plucińska et al., 2014). On the contrary, Jankowsky and co-workers showed that overexpression of mouse APP did not alter Aβ pathology when expressed on a PS1dE9 background, while it increased Aβ pathology when expressed on a more aggressive APPswe/PS1dE9 background (Jankowsky et al., 2007). These data suggest a differential effect of murine Aβ on human Aβ deposition in the different APP mouse models and may explain the mild effects observed in this study.

### Icelandic mutation and tau

Several lines of evidence suggest a connection between Aβ and tau in the pathophysiology of AD, with both proteins being abundant and often co-localising at synapses (Fein et al., 2008; Henkins et al., 2012; Kurucu et al., 2022; Sokolow et al., 2012; Tai et al., 2014). It is therefore important to quantify tau levels to confirm if they are affected by APP alterations. A study investigating the effect of the Icelandic mutation in an APP/PS1 mouse model of AD reported a decrease in phospho-tau pathology in the A673T-Aβ groups, but this reduction remains unexplained (Célestine et al., 2024). By contrast, the Icelandic A673T mutation did not affect tau levels and was unable to rescue behavioural impairment in a tau-transgenic mouse model (Anschuetz et al., 2025a). A recent exploratory study in 6 non-AD patients (unconfirmed idiopathic normal pressure hydrocephalus cases) comparing CSF of three APP^A673T^ carriers to three age- (and sex-) matched control subjects reported that disease-relevant soluble APP-β and Aβ42 levels were significantly reduced in the CSF of APP^A673T^ carries. Yet, soluble APP-α, total tau and phosphorylated tau (p-tau 181) were not altered (Wittrahm et al., 2023). This is in line with our finding that the Icelandic mutation had no effect on tau, as 5xFAD and 5xFADxAPP^A673T^ mice presented with similar tau levels. It is worth mentioning that 5xFAD showed normal tau levels not dissimilar of WT mice, and is in line with unchanged total tau levels in 5xFAD compared to WT at 3 months of age (Héraud et al., 2014).

### Icelandic mutation and synaptic proteins

Synapse loss is a key event in AD that strongly correlates with cognitive decline (de Wilde et al., 2016; Spires-Jones and Hyman, 2014). Additionally, a link between Aβ plaque formation and synaptic dysfunction has been established (Zhang et al., 2022). The presynaptic proteins SYP and SNAP25 were chosen as established markers for synapse loss in AD and AD mouse models, while STX1A was chosen as negative, non-changing, marker (Anschuetz et al., 2024, 2025b; De Wilde et al., 2016). The expression of the Icelandic mutation in 5xFAD did not alter levels of these three synaptic markers, in line with a recent report investigating the role of the Icelandic A673T mutation in a tau-based animal model (Anschuetz et al., 2025a). However, they also were unchanged across all genotypes despite previous reports of a general reduction of synaptic proteins in 5xFAD as early as 6 months (Anschuetz et al., 2025b), most notably a reduction between 30 and 45% for SYP (Jiang et al., 2020; Kim et al., 2022; Vasilopoulou et al., 2021) These discrepancies likely relate to the different quantification methods used (immunoblotting/immunofluorescence vs. ELISA).

### Icelandic mutation and neurodegeneration

Neuronal loss is a further key pathological feature of neurodegenerative disease such as AD (Crews and Masliah, 2010). Conflicting findings were reported for neuronal loss in 5xFAD mice. On one hand, stereological neuron numbers were lower in cortical layer 5 starting at 9 months (Eimer and Vassar, 2013) and persisting at 12 months (Jawhar et al., 2012), while on the other neuronal loss appeared as early as 6 months in the subiculum (Poon et al., 2023). Our analyses based on area stained in microscopic images using the ilastik software returned no significant changes of the NeuN staining in 5xFAD mice in any of the five brain regions analysed, and no effect of the Icelandic A673T mutation.

Collectively, we here show that the Icelandic mutation, A673T, has only moderate effects on Aβ pathology in 5xFAD mice, and the lack of effect is likely due to the aggressive Aβ pathology evoked at 6-month of age by the combination of the Swedish, Florida and London APP, and PS1 mutations.

## INSTITUTIONAL REVIEW BOARD STATEMENT

All animal experiments were performed in accordance with the European Communities Council Directive (63/2010/EU) and a project licence with local ethical approval under the UK Animals (Scientific Procedures) Act (1986) and its Amended Regulations (2012) and complied with the ARRIVE 2.0 guidelines. No human samples were used in this study.

## LIST OF ABBREVIATIONS

5xFAD: five familial Alzheimer’s disease mice
5xFADxAPPA673T: mouse crosses carrying both the 5xFAD mutations and the A673T mutation in the APP gene
Aβ: amyloid beta-protein
AD: Alzheimer’s disease
AEBSF: 4-(2-aminoethyl) benzenesulfonyl fluoride hydrochloride
APP: amyloid precursor protein
APPA673T: Icelandic mutation mice
BCA: bicinchoninic acid
CA1: hippocampal cornu ammonis
CB: cerebellum
CTF: carboxyl terminal fragment
CTX: visual cortex
DG: dentate gyrus
GuHCl: guanidine hydrochloride
IHC: immunohistochemistry
PFC: prefrontal cortex
PS1: presenilin-1
S.D: standard deviation
S1: RIPA: soluble supernatant fraction
S2: GuHCl fraction or RIPA-insoluble fraction
SNAP25: synaptosome associated protein 25kDa
STX1A: syntaxin 1A
SYP: synaptophysin
WT: C57Bl6/J wild-type mice

## ACKNOWLEDGEMENTS

The authors acknowledge Drs. Lianne Robinson and Valeria Melis for support with animal perfusions, and Dariia Babych for support with ELISA.

## AVAILABILITY OF DATA AND MATERIALS

Data will be made available on reasonable request.

## AUTHORS’ CONTRIBUTIONS

AA: Investigation, Data curation, Formal analysis, Visualization, Writing - Original Draft; RL: Investigation; TV: Investigation; BP: Conceptualization, Resources, Writing - Review & Editing; CRH: Funding acquisition; GR: Conceptualization, Supervision, Project administration, Funding acquisition, Writing - Review & Editing; KS: Conceptualization, Supervision, Investigation, Data curation, Formal analysis, Visualization, Writing - Original Draft.

---

**Figure S1:**
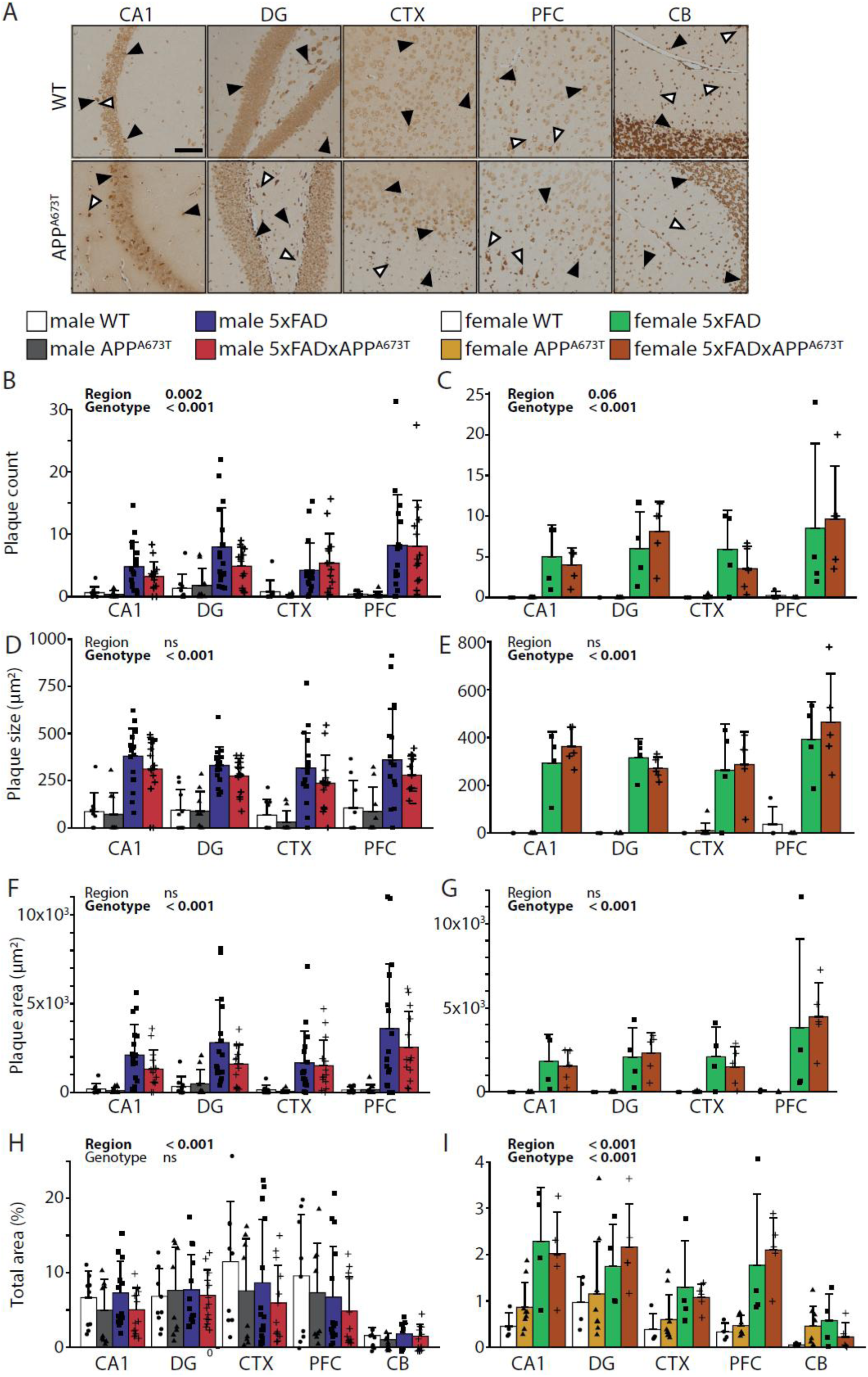
Aβ immunohistochemistry using the antibody 6E10. (A) Representative Aβ immunohistochemistry images in brains of male WT and APP^A673T^ mice stained with the antibody 6E10 (Biolegend # 803004, diluted 1:1,000). For the other two genotypes (5xFAD and 5xFADxAPP^A673T^) see Fig. 1, main manuscript. Images from CA1, DG, CTX and PFC were taken using a light microscope at a 100x magnification. *Black arrowheads, cytosolic staining; white arrowheads, axonal/dendritic staining; scale bars, 100µm*. Aβ plaque levels were quantified using ilastik as plaque counts (B&C), plaque size (D&E), plaque area (F&G) and total area stained (H&I) in male (B, D, F and H) and female (C, E, G and I) WT, APP^A673T^, 5xFAD and 5xFADxAPP^A673T^ mice in four pre-specified brain regions. Data is shown as individual values, group mean, and S.D. Data were analysed using Wald χ² test (B,C) or two-way ANOVA (D-I) with genotype and region as independent variables. Count data did not need to be transformed (B,C) and remaining data were transformed using Yeo-Johnson (D-G) or square-root (H, I) transformation. Males: WT: n=9, A673T: n=9, 5xFAD: n=17 (PFC n=16), 5xFADxAPP^A673T^: n=14 (PFC n=13). Females: WT: n=4 (CB n=3), A673T: n=9 (PFC n=6, CB n=4), 5xFAD: n=4, 5xFADxAPP^A673T^: n=5. Abbreviations: CA1: hippocampal CA1, CTX: visual cortex, DG: dentate gyrus, ns: not significant, PFC: prefrontal cortex

**Figure S2:**
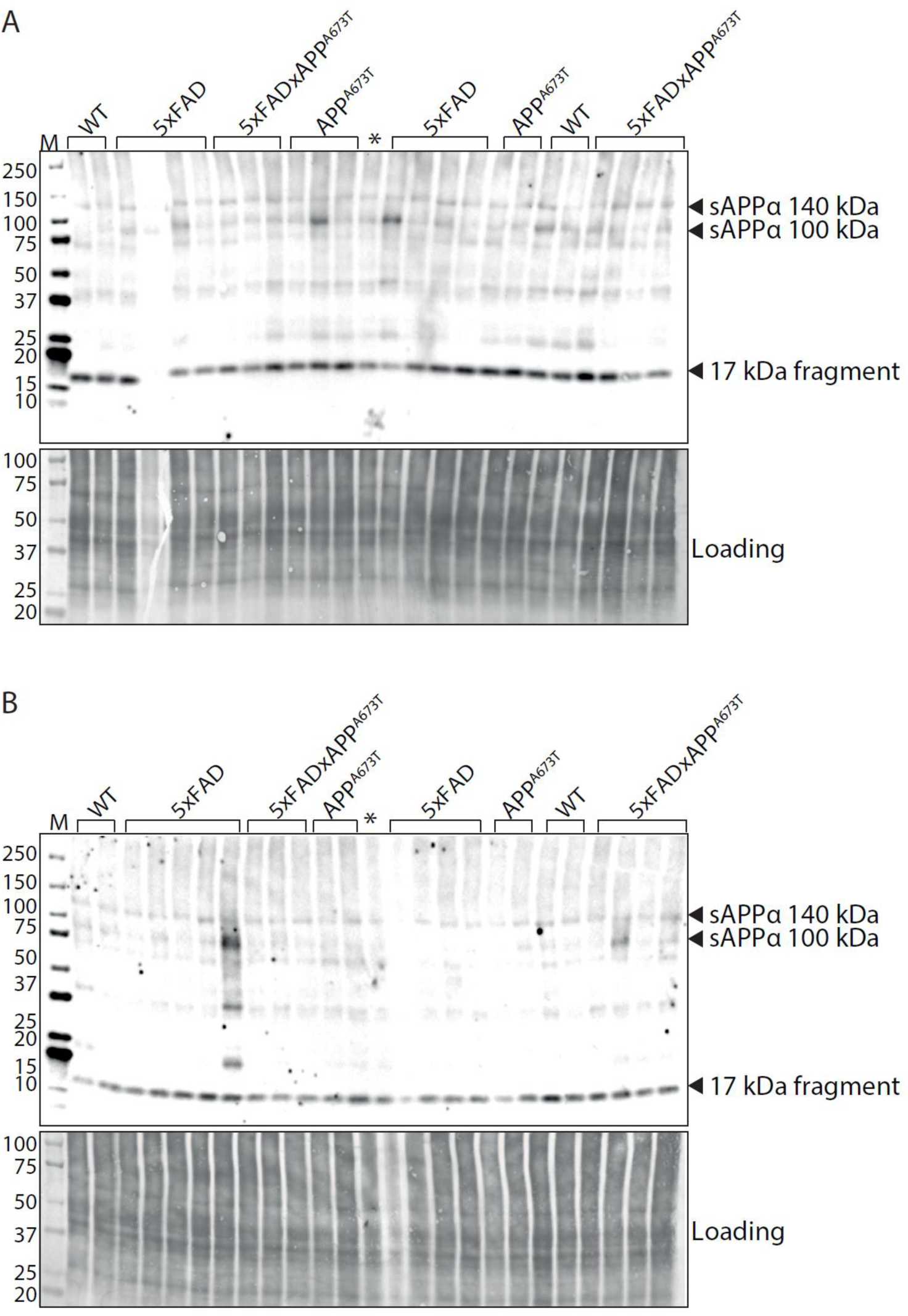
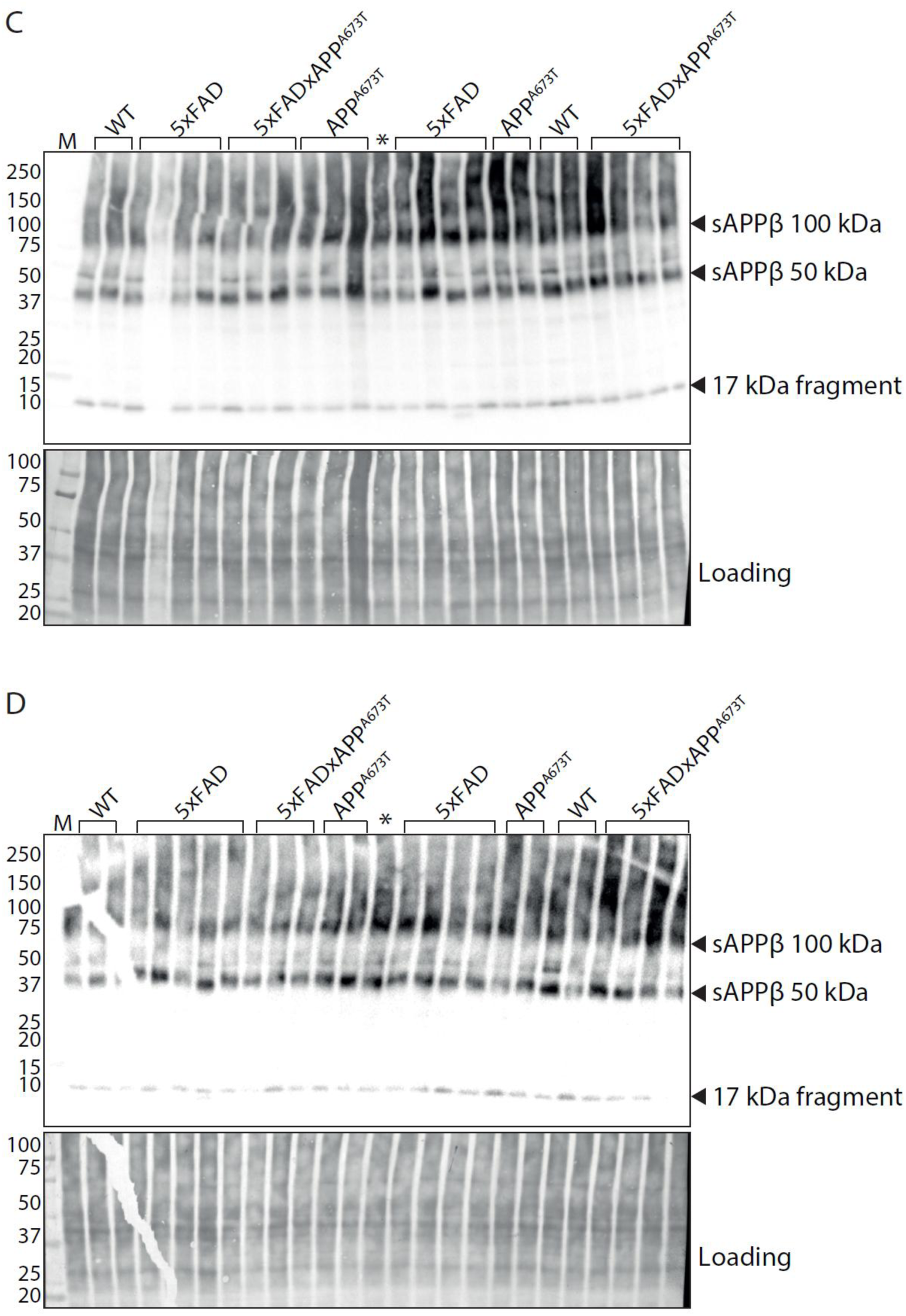
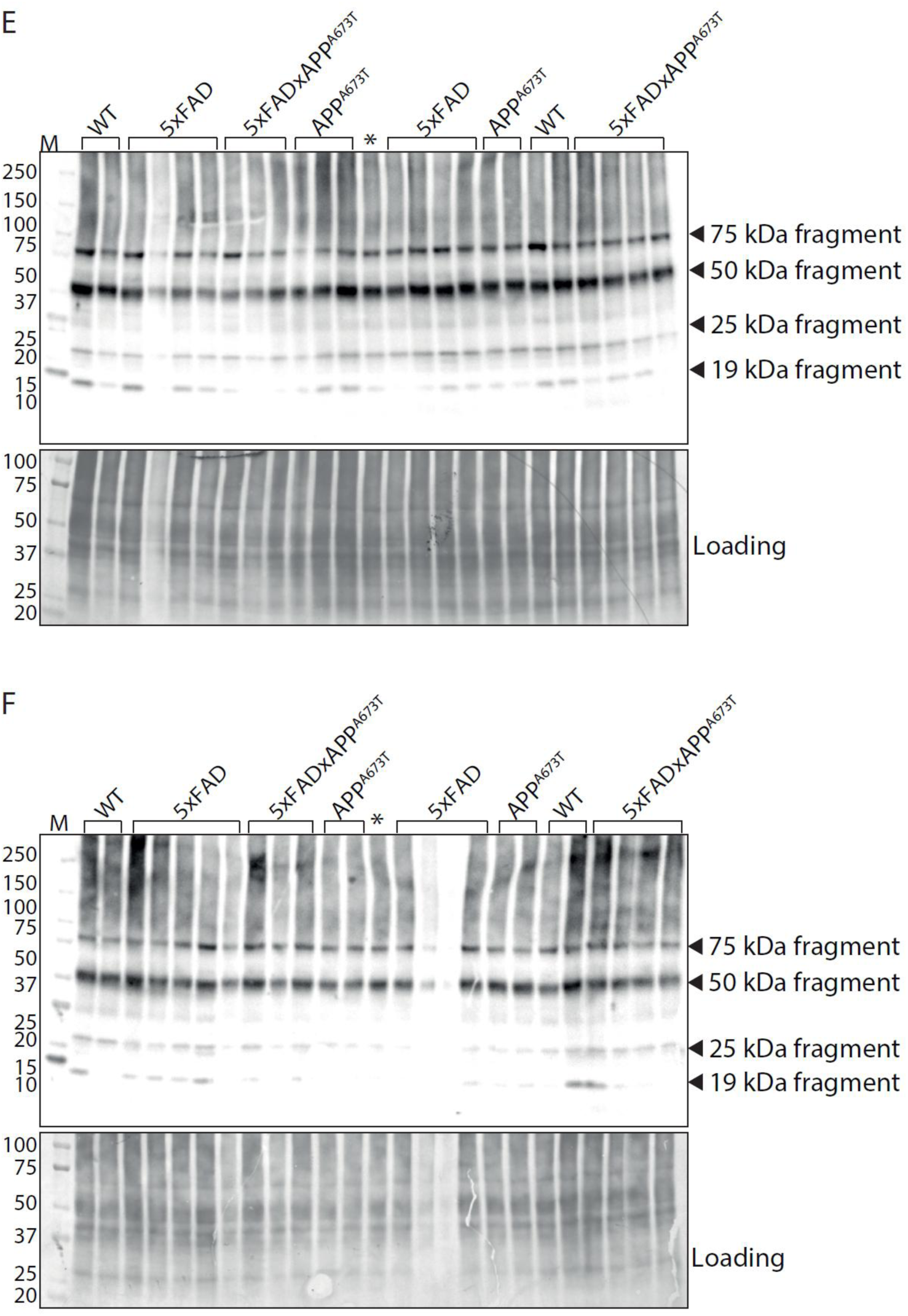
Quantification of APP/Aβ species using immunoblotting. Proteins from RIPA-soluble S1 fractions were separated by SDS-PAGE (20 µg per lane) and labelled with the antibodies 2B3 (IBL #11088, 1:500) against sAPPα (A&B), Poly8134 (Biolegend #Poly8134, 1:1,000) against sAPPβ (C&D), CT695 (Invitrogen #51-2700, 1:1,000) against CTFs of APP (E&F), and all blots for male WT, APP^A673T^, 5xFAD and 5xFADxAPP^A673T^ mice are displayed. For densitometric quantification see Fig. 3, main text. WT: n=8 (n=6 for Poly8134 antibody), A673T: n=9, 5xFAD: n=17 (n=15 for Poly8134 antibody), 5xFADx APP^A673T^: n=14.

**Figure S3:**
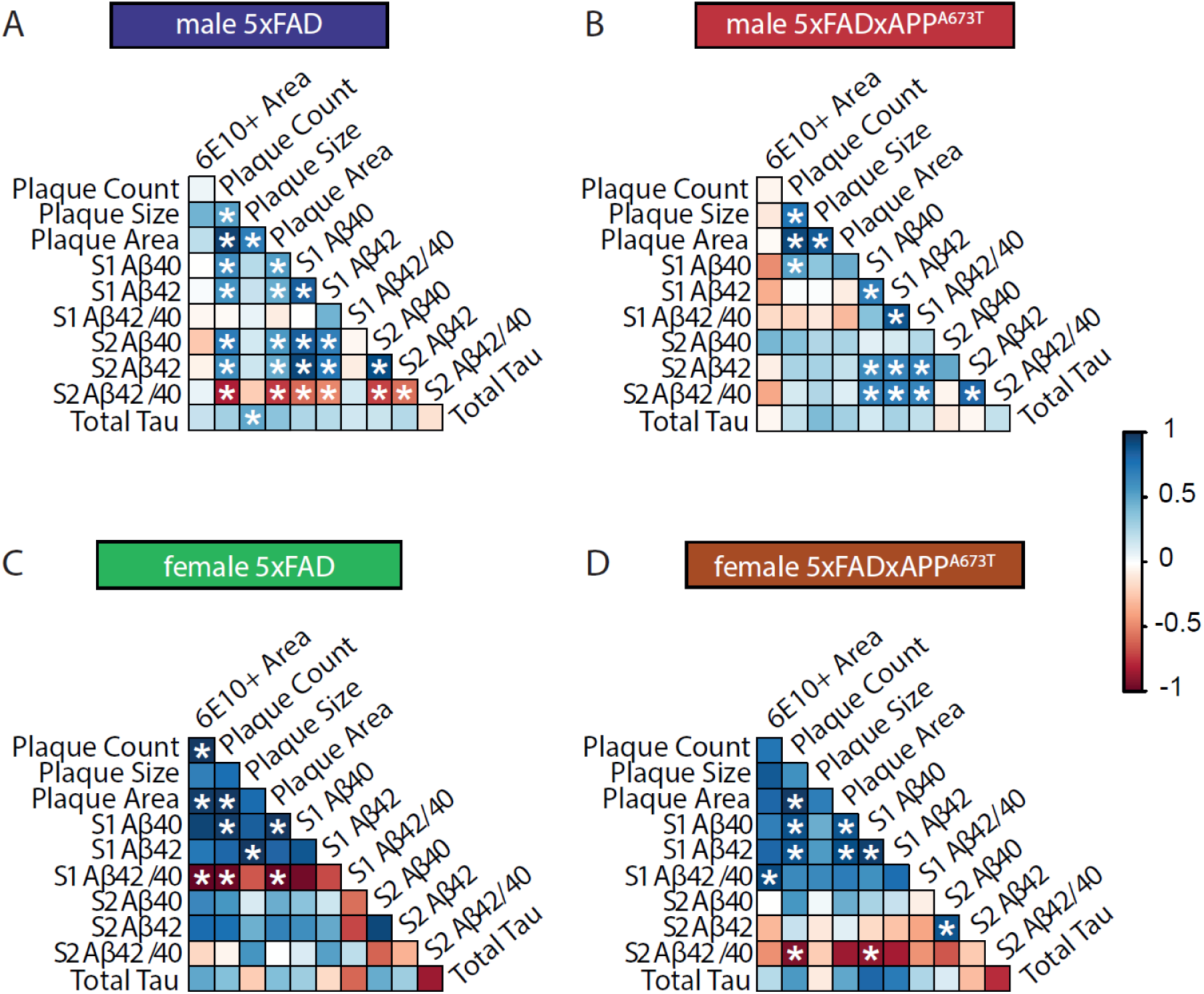
Correlation matrices for the different Aβ measurements and tau. (A-D) Pearson correlation matrix between Aβ plaque parameters (count, size, and area), Aβ40 and Aβ42 levels and their ratio (Aβ42/ Aβ40) in soluble and insoluble fractions S1 and S2, as well as tau ELISA are displayed for 5xFAD male (A), 5xFADxAPP^A673T^ male (B), 5xFAD female (C) and 5xFADxAPP^A673T^ female (D) mice with blue for positive correlations, red for negative correlations and white where no correlation was seen (* p < 0.05). Aβ and tau levels were quantified using ELISA and plaque counts, size and total area were quantified using immunohistochemistry (averaged across brain regions). Data were analysed using Jennrich test to detect differences between matrices. Abbreviations: APPA673T: Icelandic mutation mice, 5xFAD: five familial Alzheimer’s disease mice, 5xFADxAPP^A673T^: crosses carrying both the 5xFAD mutations and the A673T mutation in the APP gene.

**Figure S4:**
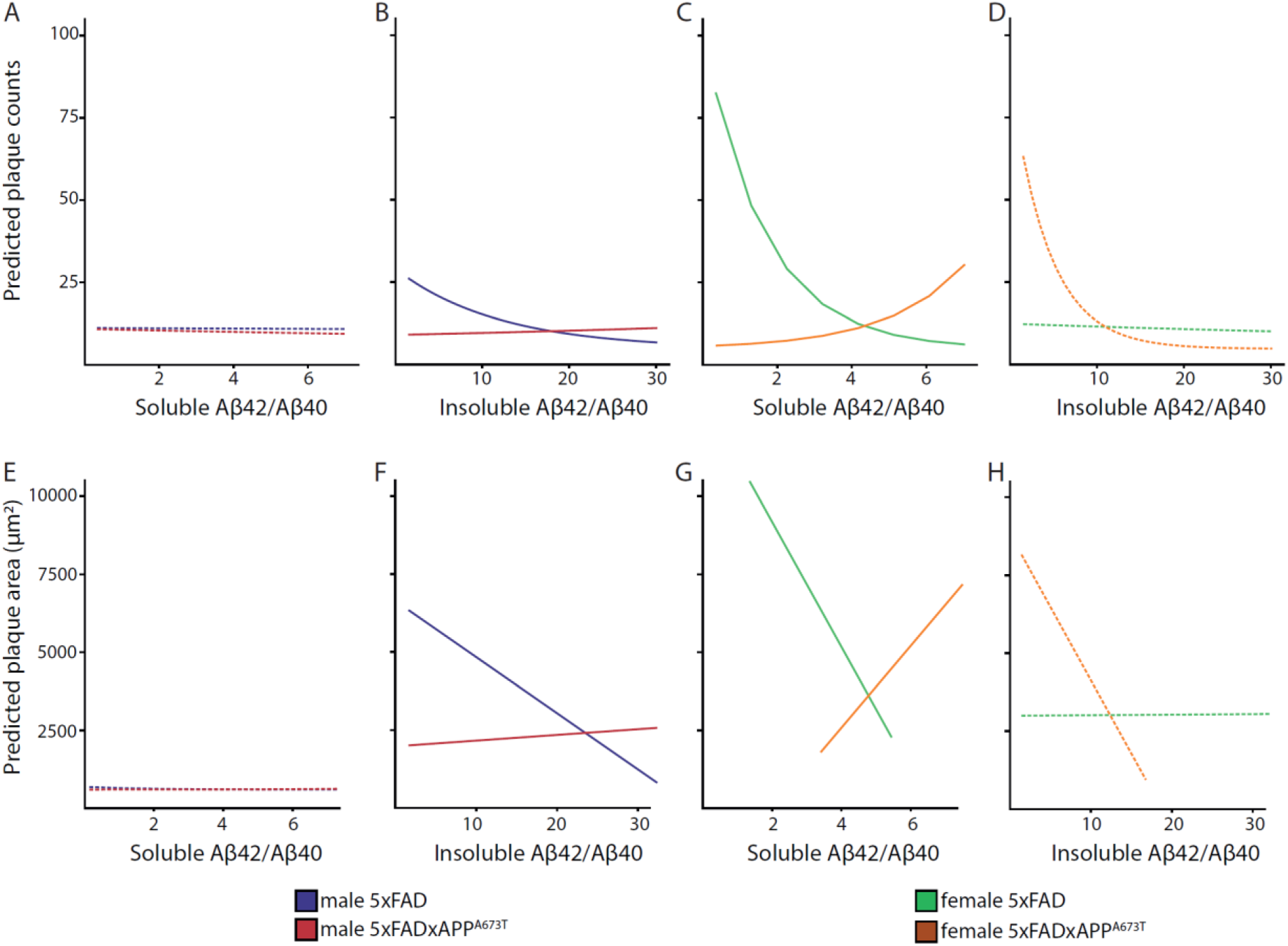
Prediction linear modelling for the different Aβ measurements. Generalized linear modelling was used to explore the effect of Aβ42/Aβ42 ratio in S1 and S2 on plaque counts (A-D) and plaque area (E-H)), and whether this differed across genotypes. Model-predicted plaque counts depending on Aβ42/Aβ40 ratio for 5xFAD and 5xFADxAPP^A673T^ male in S1 (A) and S2 (B), as well as 5xFAD and 5xFADxAPP^A673T^ female in S1 (C) and S2 (D) are presented. Further, model-predicted plaque area depending on Aβ42/Aβ40 ratio for 5xFAD and 5xFADxAPP^A673T^ male in S1 (E) and S2 (F), as well as 5xFAD and 5xFADxAPP^A673T^ female in S1 (G) and S2 (H) are presented. Dashed lines indicate non-significant effects.

